# Emergence of an Antigenically Drifted and Reassorted Influenza B Virus at the end of the 2024-25 Influenza Season

**DOI:** 10.1101/2025.07.24.666632

**Authors:** Elgin Akin, David A. Villafuerte, Anne P. Werner, Matthew Pinsley, Corinne Pierce, Amary Fall, Omar Abdullah, Julie M. Norton, Richard E. Rothman, Katherine Z.J. Fenstermacher, Yu-Nong Gong, Eili Klein, Heba H. Mostafa, Andrew Pekosz

## Abstract

Influenza B virus (IBV) is a significant contributor to annual and severe cases of influenza, particularly in the young and elderly. Late in the 2024-25 Northern Hemisphere influenza season, a surge of IBV cases were identified in the Johns Hopkins Hospital Systems. The IBV responsible for the surge, C.3.1/re, was a clade C.3 virus that had reassorted with clade C.5.1 viruses and acquired the D197N mutation in hemagglutinin restoring a putative N-linked glycan predicted to mask a key neutralizing antibody epitope. The C.3.1/re viruses preferentially infected children but showed no significant change in disease severity. C.3.1/re viruses were poorly neutralized by pre- and post- influenza vaccination serum in a human cohort. Removal of the glycan at residue 197 restored neutralizing antibody recognition. The C.3.1/re IBV genotype that emerged late in the 2024-25 influenza season was antigenically mismatched with IBV vaccine strains for the 2025 and 2026 Southern hemisphere, as well as the 2025-26 Northern Hemisphere influenza seasons. While the 2026-27 Northern Hemisphere vaccine strain is a C.3.1/re, the egg adapted isolate selected (B/Tokyo/EIS13-175/2025) lacks the 197 glycosylation which is predicted to have poor recognition with circulating IBV clades. Phylogenetic analysis of currently circulating IBVs shows a diversification of circulating C.3 clades with multiple reassortment events between C.3 and C.5 clades in addition to independent acquisitions of D197N mutations, suggesting IBV is going through a period of significant antigenic and genetic expansion.

**IMPORTANCE:** Influenza B viruses are undergoing a period of antigenic and genetic expansion, with several reassorted viruses emerging that also contain point mutations in key hemagglutinin antigenic sites proximal to the receptor binding domain. This has important impacts on vaccine strain choice, as only one IBV component is included in current influenza vaccines. We demonstrate a significant shift in the demographics of IBV infected individuals with the emergence of the antigenically drifted and reassorted IBV C.3.1/re. Furthermore, we show that 197 glycosylation of hemagglutinin is critical for C.3.1/re antigenic drift and we document several emergent C.3 reassortments encoding the D197N mutation. With the IBV vaccine component for the Northern Hemisphere 2026-27 season having lost a key N-linked glycan on the hemagglutinin protein, and multiple independent emergences of antigenically drifted and reassorted viruses, attention to IBV infections should be increased in the upcoming Southern and Northern hemisphere influenza seasons.

## INTRODUCTION

Seasonal influenza epidemics, caused by influenza A and B viruses, remain a major global health burden, resulting in an estimated 250,000–500,000 deaths annually. Although influenza A viruses have received greater attention due to their pandemic potential, influenza B viruses (IBV) contribute substantially to seasonal morbidity and mortality, particularly among pediatric populations ^1–3^. Although IBV evolves more slowly than influenza A virus (IAV), leading to less frequent antigenic drift and extended geographic persistence, it remains capable of driving significant outbreaks and exhibits variable vaccine effectiveness, underscoring the need to further define mechanisms of antigenic evolution^4^. The disproportionate impact of IBV on pediatric populations, combined with evidence of reduced vaccine effectiveness against IBV in certain seasons, underscores the need for vigilant molecular surveillance of circulating strains.

Antigenic drift in IBV is primarily mediated by mutations in the hemagglutinin (HA) glycoprotein which enables receptor binding and membrane fusion and is the principal target of neutralizing antibodies ^5–9^. Furthermore. mutations in HA can impact receptor binding ^6–8,10^, while mutations in other gene segments can alter replication efficiency ^11–13^, and innate immune evasion^14–16^. If two genetically distinct viruses infect the same host cell, the segmented nature of the genome also allows for reassortment, resulting in progeny with novel constellations of gene segments. Such reassortments can affect antigenic properties, virus replication dynamics or antiviral drug sensitivity ^17–20^.

Beyond amino acid substitutions, the gain or loss of N-linked glycans can profoundly alter antigenicity by masking epitopes, particularly near the receptor binding site (RBS) in the HA protein ^9,21^. The 190-helix is a structurally constrained region that contributes to receptor engagement and is a known target of neutralizing antibodies, making it highly sensitive to such modifications ^5,22,23^. In IBVs, an asparagine at HA residue 197 (B/Brisbane/06/2008 numbering) has been conserved since the emergence of the B/Victoria lineage, encoding an N-linked glycosylation site proximal to the RBS ^24^. Loss of this glycan has been repeatedly observed during egg adaptation, enhancing viral growth in embryonated chicken eggs and altering antigenic identity ^25–27^. While the antigenic consequences of glycan loss at residue 197 have been characterized in the role of egg-adapted vaccine strains using ferret sera, the effects of glycan gain or restoration at this site during natural human circulation, and more broadly the frequency and immunological impact of glycosylation changes near the 190-helix in circulating IBV populations, remain poorly defined ^26^.

During the 2024–25 Northern Hemisphere influenza season, a late-season surge of IBV cases within the Johns Hopkins Hospital System coincided with a shift in circulating IBV subclades. This increase was driven by the rapid expansion of C.3.1, a C.3-derived lineage. Notably, C.3 had persisted at low levels since its initial detection in 2023, whereas the previously dominant C.5.1 subclade and other C.5 descendants, C.5.6 and C.5.7, were detected at unexpectedly low frequencies, motivating investigation into the molecular basis of C.3.1 emergence. C.3.1/re is defined by putative reassortment with C.5.1 and a HA1:D197N reversion that restores a glycosylation motif within the 190-helix, together with a secondary HA1:P208S substitution. It is hypothesized that the 197 glycan may alter epitope accessibility and reduce susceptibility to vaccine-induced neutralizing antibodies. To investigate this, genomic surveillance, phylogenetic reconstruction, infectious IBV clones and serology were integrated to characterize the evolutionary origins and antigenic properties of C.3.1. These data indicate that the reversion of glycosylation at HA residue 197 is associated with reduced neutralization by vaccine-induced sera, consistent with a potential contribution to antigenic drift.

## RESULTS

### Replacement of the Influenza B virus subclade C.5.1 with C.3.1 in the 2024-25 season

Within the Johns Hopkins Hospital System (JHHS), 15% (894/5365) of influenza-positive patients were infected with influenza B virus (IBV) in the 2024-25 season **(Table S1).** IBV cases rose sharply from 65 total cases in February to 372 and 400 cases in March and April respectively **(Fig. 1A and Table S1).** A subset of IBV+ specimens were subject to whole genome sequencing and 58.5% belonged to the C.3 subclade **(Fig. 1B**, **Table 1)**. Subclade C.5.6 was detected at 22.3% and the formerly dominant 2023-24 subclade C.5.1 at 11.5% **(Table 1)**. C.5.7 and the parental C.5 subclades were detected in comparatively low abundance at 3.8% and 3.1%, respectively. Nationally, the percentage of C.3 infections was much lower than what was observed in the state of Maryland (18.3 versus 58.5%, **Table 1**).

**Figure 1.**
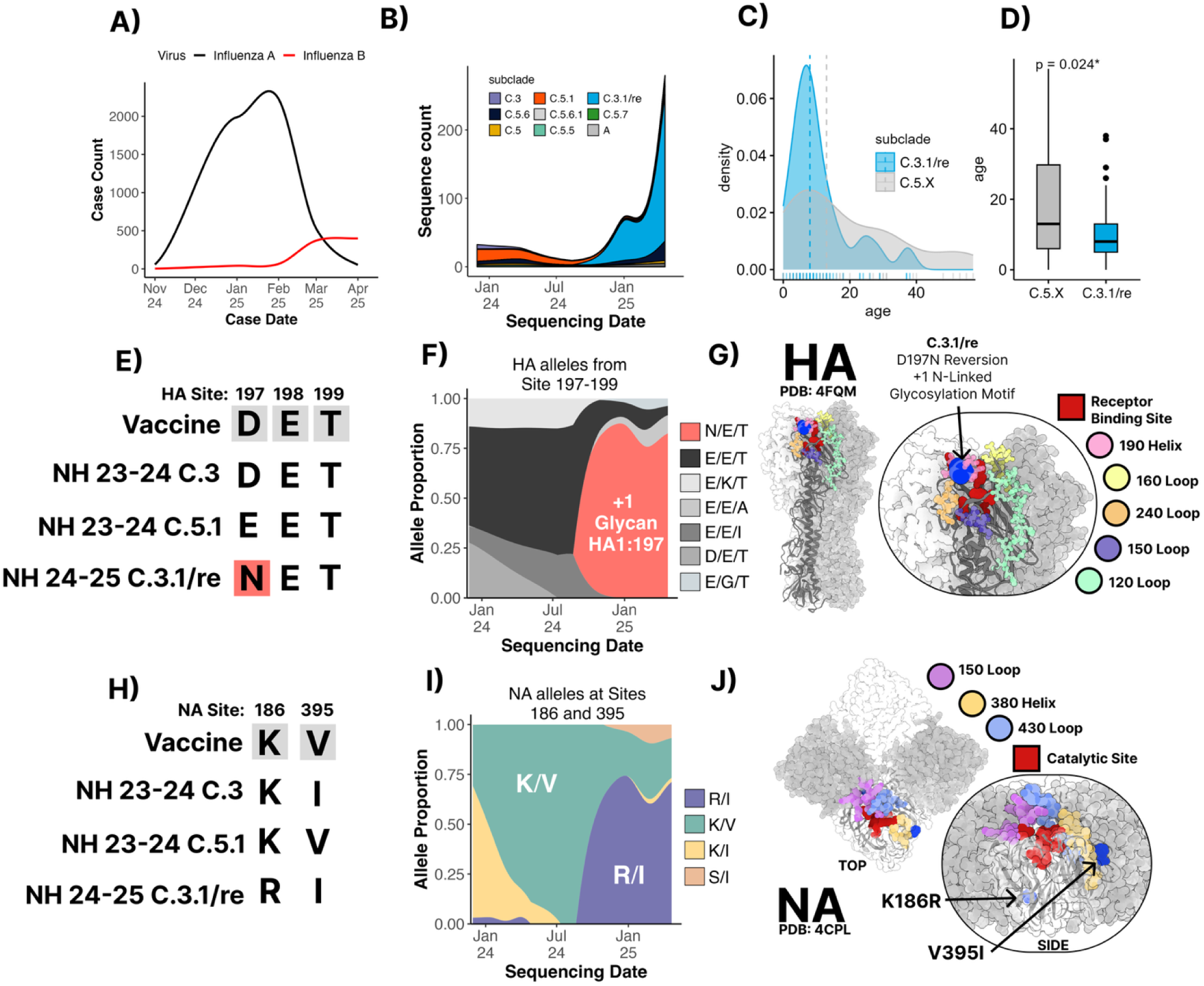
Emergence of Influenza B/Victoria/16/1987-like C.3.1 subclade, acquisition of a novel glycosylation site, and age-related infection bias. **(A)** Weekly clinical case counts of influenza A (black) and B (red) reported from November 2024 to April 2025. **(B)** Longitudinal distribution of IBV B/Victoria HA subclades (colored by clade) based on sequencing data. **(C)** Kernel density estimate of patient age for sequences classified as C.3 (blue) or C.5.X (gray). Tick marks below each curve indicate individual ages. **(D)** Boxplot comparison of age distribution for C.3 versus all C.5-lineages (C.5.X) (Wilcoxon test, *p* = 0.024). **(E)** Sequence alignment of HA (residues 197–199) protein motifs across the vaccine strain, parental C.5.1, C.3 and 2024-25 C.3 (B/Brisbane/08/2008 numbering). **(F)** The 2024–25 C.3 subclade acquired an N-linked glycosylation site with a 197N mutation, whereas C.5.1 retained the 197E motif. **(G)** Structural mapping of the D197N substitution onto the HA trimer (PDB: 4FMQ). Influenza B HA Antigenic sites are annotated. **(H)** NA sequence alignments reveal substitutions V395I and K186R. **(I)** the penetrance of mutations at positions 395 and 186 in the population over time. **(J)** NA structures showing the position of the V395I substitution in the 380-helix antigenic site and the K186R on the basal side of the NA enzymatic active site.

**Table 1.** 2024-25 Influenza B virus subclade sequence counts submitted by the Johns Hopkins Hospital System (JHHS) or uploaded to GISAID filtered to North American sequences.

| Subclade | Global | Johns Hopkins Hospital System (JHHS) | North America - Excluding JHHS |
| --- | --- | --- | --- |
| C.2 | 11 (0.5%) | 0 (0%) | 0 (0%) |
| C.3 | 5 (0.2%) | 0 (0%) | 0 (0%) |
| C.3.1 | 2 (0.1%) | 75 (58.1%) | 115 (20.5%) |
| C.3.2 | 1 (0%) | 1 (0.8%) | 10 (1.8%) |
| C.5 | 35 (1.7%) | 4 (3.1%) | 56 (10%) |
| C.5.1 | 729 (36.3%) | 15 (11.6%) | 193 (34.3%) |
| C.5.6 | 511 (25.4%) | 27 (20.9%) | 83 (14.8%) |
| C.5.6.1 | 238 (11.8%) | 2 (1.6%) | 12 (2.1%) |
| C.5.7 | 478 (23.8%) | 5 (3.9%) | 93 (16.5%) |

Data Source: JHHS and GISAID between 2024-10-01 to 2025-05-31

**Table 2.**
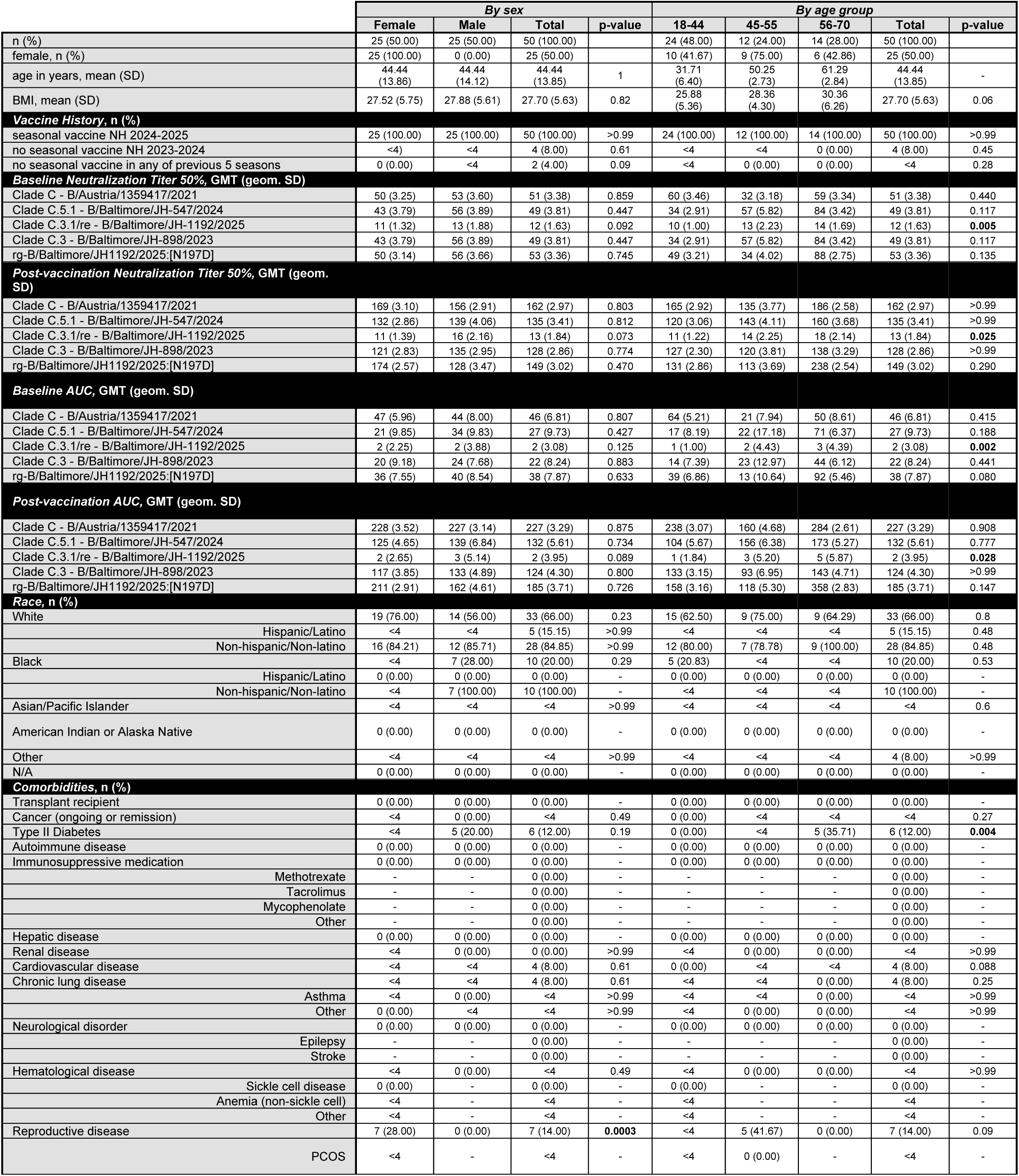
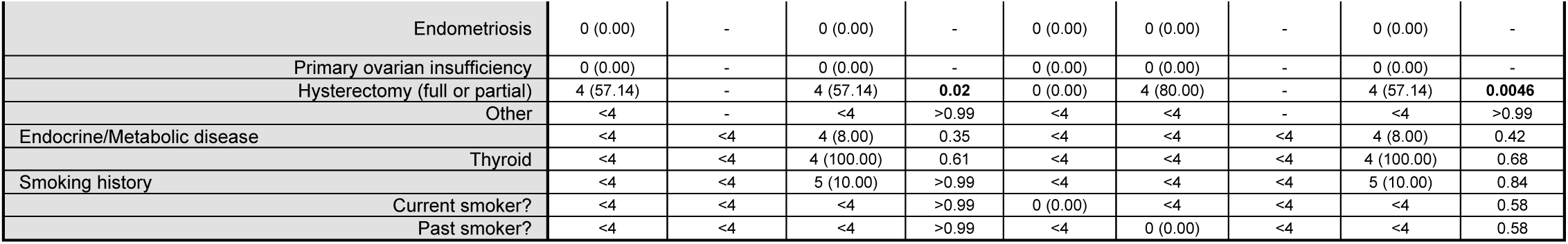
2024-25 JHJ-CEIRR Vaccine Cohort characteristics, demographics, and geometric mean titers for all assessments described. Per Internal Review Board (IRB) directive, categories representing less than 4 individuals in total are summarized as <4. For qualitative data and co-morbidities, Fisher’s exact test was used to generate p-values shown. For baseline and post-vaccination immunity readouts, adjusted p-values shown were calculated via Kruskal-Wallis nonparametric test with Bonferroni’s correction for age analyses, and Dunn’s test for multiple comparisons for sex analyses, both with Bonferroni’s corrections. Bolded p-values are significant (i.e., p < 0.05).

|  | By sex |  |  |  | By age group |  |  |  |  |
| --- | --- | --- | --- | --- | --- | --- | --- | --- | --- |
|  | Female | Male | Total | p-value | 18-44 | 45-55 | 56-70 | Total | p-value |
| n (%) | 25 (50.00) | 25 (50.00) | 50 (100.00) |  | 24 (48.00) | 12 (24.00) | 14 (28.00) | 50 (100.00) |  |
| female, n (%) | 25 (100.00) | 0 (0.00) | 25 (50.00) |  | 10 (41.67) | 9 (75.00) | 6 (42.86) | 25 (50.00) |  |
| age in years, mean (SD) | 44.44<br>(13.86) | 44.44<br>(14.12) | 44.44<br>(13.85) | 1 | 31.71<br>(6.40) | 50.25<br>(2.73) | 61.29<br>(2.84) | 44.44<br>(13.85) | - |
| BMI, mean (SD) | 27.52 (5.75) | 27.88 (5.61) | 27.70 (5.63) | 0.82 | 25.88<br>(5.36) | 28.36<br>(4.30) | 30.36<br>(6.26) | 27.70 (5.63) | 0.06 |
| **Vaccine History, n (%)** |  |  |  |  |  |  |  |  |  |
| seasonal vaccine NH 2024-2025 | 25 (100.00) | 25 (100.00) | 50 (100.00) | >0.99 | 24 (100.00) | 12 (100.00) | 14 (100.00) | 50 (100.00) | >0.99 |
| no seasonal vaccine NH 2023-2024 | <4 | <4 | 4 (8.00) | 0.61 | <4 | <4 | 0 (0.00) | 4 (8.00) | 0.45 |
| no seasonal vaccine in any of previous 5 seasons | 0 (0.00) | <4 | 2 (4.00) | 0.09 | <4 | 0 (0.00) | 0 (0.00) | <4 | 0.28 |
| **Baseline Neutralization Titer 50%, GMT (geom. SD)** |  |  |  |  |  |  |  |  |  |
| Clade C - B/Austria/1359417/2021 | 50 (3.25) | 53 (3.60) | 51 (3.38) | 0.859 | 60 (3.46) | 32 (3.18) | 59 (3.34) | 51 (3.38) | 0.440 |
| Clade C.5.1 - B/Baltimore/JH-547/2024 | 43 (3.79) | 56 (3.89) | 49 (3.81) | 0.447 | 34 (2.91) | 57 (5.82) | 84 (3.42) | 49 (3.81) | 0.117 |
| Clade C.3.1/re - B/Baltimore/JH-1192/2025 | 11 (1.32) | 13 (1.88) | 12 (1.63) | 0.092 | 10 (1.00) | 13 (2.23) | 14 (1.69) | 12 (1.63) | **0.005** |
| Clade C.3 - B/Baltimore/JH-898/2023 | 43 (3.79) | 56 (3.89) | 49 (3.81) | 0.447 | 34 (2.91) | 57 (5.82) | 84 (3.42) | 49 (3.81) | 0.117 |
| rg-B/Baltimore/JH1192/2025:[N197D] | 50 (3.14) | 56 (3.66) | 53 (3.36) | 0.745 | 49 (3.21) | 34 (4.02) | 88 (2.75) | 53 (3.36) | 0.135 |
| **Post-vaccination Neutralization Titer 50%, GMT (geom. SD)** |  |  |  |  |  |  |  |  |  |
| Clade C - B/Austria/1359417/2021 | 169 (3.10) | 156 (2.91) | 162 (2.97) | 0.803 | 165 (2.92) | 135 (3.77) | 186 (2.58) | 162 (2.97) | >0.99 |
| Clade C.5.1 - B/Baltimore/JH-547/2024 | 132 (2.86) | 139 (4.06) | 135 (3.41) | 0.812 | 120 (3.06) | 143 (4.11) | 160 (3.68) | 135 (3.41) | >0.99 |
| Clade C.3.1/re - B/Baltimore/JH-1192/2025 | 11 (1.39) | 16 (2.16) | 13 (1.84) | 0.073 | 11 (1.22) | 14 (2.25) | 18 (2.14) | 13 (1.84) | **0.025** |
| Clade C.3 - B/Baltimore/JH-898/2023 | 121 (2.83) | 135 (2.95) | 128 (2.86) | 0.774 | 127 (2.30) | 120 (3.81) | 138 (3.29) | 128 (2.86) | >0.99 |
| rg-B/Baltimore/JH1192/2025:[N197D] | 174 (2.57) | 128 (3.47) | 149 (3.02) | 0.470 | 131 (2.86) | 113 (3.69) | 238 (2.54) | 149 (3.02) | 0.290 |
| **Baseline AUC, GMT (geom. SD)** |  |  |  |  |  |  |  |  |  |
| Clade C - B/Austria/1359417/2021 | 47 (5.96) | 44 (8.00) | 46 (6.81) | 0.807 | 64 (5.21) | 21 (7.94) | 50 (8.61) | 46 (6.81) | 0.415 |
| Clade C.5.1 - B/Baltimore/JH-547/2024 | 21 (9.85) | 34 (9.83) | 27 (9.73) | 0.427 | 17 (8.19) | 22 (17.18) | 71 (6.37) | 27 (9.73) | 0.188 |
| Clade C.3.1/re - B/Baltimore/JH-1192/2025 | 2 (2.25) | 2 (3.88) | 2 (3.08) | 0.125 | 1 (1.00) | 2 (4.43) | 3 (4.39) | 2 (3.08) | **0.002** |
| Clade C.3 - B/Baltimore/JH-898/2023 | 20 (9.18) | 24 (7.68) | 22 (8.24) | 0.883 | 14 (7.39) | 23 (12.97) | 44 (6.12) | 22 (8.24) | 0.441 |
| rg-B/Baltimore/JH1192/2025:[N197D] | 36 (7.55) | 40 (8.54) | 38 (7.87) | 0.633 | 39 (6.86) | 13 (10.64) | 92 (5.46) | 38 (7.87) | 0.080 |
| **Post-vaccination AUC, GMT (geom. SD)** |  |  |  |  |  |  |  |  |  |
| Clade C - B/Austria/1359417/2021 | 228 (3.52) | 227 (3.14) | 227 (3.29) | 0.875 | 238 (3.07) | 160 (4.68) | 284 (2.61) | 227 (3.29) | 0.908 |
| Clade C.5.1 - B/Baltimore/JH-547/2024 | 125 (4.65) | 139 (6.84) | 132 (5.61) | 0.734 | 104 (5.67) | 156 (6.38) | 173 (5.27) | 132 (5.61) | 0.777 |
| Clade C.3.1/re - B/Baltimore/JH-1192/2025 | 2 (2.65) | 3 (5.14) | 2 (3.95) | 0.089 | 1 (1.84) | 3 (5.20) | 5 (5.87) | 2 (3.95) | **0.028** |
| Clade C.3 - B/Baltimore/JH-898/2023 | 117 (3.85) | 133 (4.89) | 124 (4.30) | 0.800 | 133 (3.15) | 93 (6.95) | 143 (4.71) | 124 (4.30) | >0.99 |
| rg-B/Baltimore/JH1192/2025:[N197D] | 211 (2.91) | 162 (4.61) | 185 (3.71) | 0.726 | 158 (3.16) | 118 (5.30) | 358 (2.83) | 185 (3.71) | 0.147 |
| **Race, n (%)** |  |  |  |  |  |  |  |  |  |
| White | 19 (76.00) | 14 (56.00) | 33 (66.00) | 0.23 | 15 (62.50) | 9 (75.00) | 9 (64.29) | 33 (66.00) | 0.8 |
| Hispanic/Latino | <4 | <4 | 5 (15.15) | >0.99 | <4 | <4 | <4 | 5 (15.15) | 0.48 |
| Non-hispanic/Non-latino | 16 (84.21) | 12 (85.71) | 28 (84.85) | >0.99 | 12 (80.00) | 7 (78.78) | 9 (100.00) | 28 (84.85) | 0.48 |
| Black | <4 | 7 (28.00) | 10 (20.00) | 0.29 | 5 (20.83) | <4 | <4 | 10 (20.00) | 0.53 |
| Hispanic/Latino | 0 (0.00) | 0 (0.00) | 0 (0.00) | - | 0 (0.00) | 0 (0.00) | 0 (0.00) | 0 (0.00) | - |
| Non-hispanic/Non-latino | <4 | 7 (100.00) | 10 (100.00) | - | <4 | <4 | <4 | 10 (100.00) | - |
| Asian/Pacific Islander | <4 | <4 | <4 | >0.99 | <4 | <4 | <4 | <4 | 0.6 |
| American Indian or Alaska Native | 0 (0.00) | 0 (0.00) | 0 (0.00) | - | 0 (0.00) | 0 (0.00) | 0 (0.00) | 0 (0.00) | - |
| Other | <4 | <4 | <4 | >0.99 | <4 | <4 | <4 | 4 (8.00) | >0.99 |
| N/A | 0 (0.00) | 0 (0.00) | 0 (0.00) | - | 0 (0.00) | 0 (0.00) | 0 (0.00) | 0 (0.00) | - |
| **Comorbidities, n (%)** |  |  |  |  |  |  |  |  |  |
| Transplant recipient | 0 (0.00) | 0 (0.00) | 0 (0.00) | - | 0 (0.00) | 0 (0.00) | 0 (0.00) | 0 (0.00) | - |
| Cancer (ongoing or remission) | <4 | 0 (0.00) | <4 | 0.49 | 0 (0.00) | <4 | <4 | <4 | 0.27 |
| Type II Diabetes | <4 | 5 (20.00) | 6 (12.00) | 0.19 | 0 (0.00) | <4 | 5 (35.71) | 6 (12.00) | **0.004** |
| Autoimmune disease | 0 (0.00) | 0 (0.00) | 0 (0.00) | - | 0 (0.00) | 0 (0.00) | 0 (0.00) | 0 (0.00) | - |
| Immunosuppressive medication | 0 (0.00) | 0 (0.00) | 0 (0.00) | - | 0 (0.00) | 0 (0.00) | 0 (0.00) | 0 (0.00) | - |
| Methotrexate | - | - | 0 (0.00) | - | - | - | - | 0 (0.00) | - |
| Tacrolimus | - | - | 0 (0.00) | - | - | - | - | 0 (0.00) | - |
| Mycophenolate | - | - | 0 (0.00) | - | - | - | - | 0 (0.00) | - |
| Other | - | - | 0 (0.00) | - | - | - | - | 0 (0.00) | - |
| Hepatic disease | 0 (0.00) | 0 (0.00) | 0 (0.00) | - | 0 (0.00) | 0 (0.00) | 0 (0.00) | 0 (0.00) | - |
| Renal disease | <4 | 0 (0.00) | 0 (0.00) | >0.99 | <4 | 0 (0.00) | 0 (0.00) | <4 | >0.99 |
| Cardiovascular disease | <4 | <4 | 4 (8.00) | 0.61 | 0 (0.00) | <4 | <4 | 4 (8.00) | 0.088 |
| Chronic lung disease | <4 | <4 | 4 (8.00) | 0.61 | <4 | <4 | 0 (0.00) | 4 (8.00) | 0.25 |
| Asthma | <4 | 0 (0.00) | <4 | >0.99 | <4 | <4 | 0 (0.00) | <4 | >0.99 |
| Other | 0 (0.00) | <4 | <4 | >0.99 | <4 | 0 (0.00) | 0 (0.00) | <4 | >0.99 |
| Neurological disorder | 0 (0.00) | 0 (0.00) | 0 (0.00) | - | 0 (0.00) | 0 (0.00) | 0 (0.00) | 0 (0.00) | - |
| Epilepsy | - | - | 0 (0.00) | - | - | - | - | 0 (0.00) | - |
| Stroke | - | - | 0 (0.00) | - | - | - | - | 0 (0.00) | - |
| Hematological disease | <4 | 0 (0.00) | <4 | 0.49 | <4 | 0 (0.00) | 0 (0.00) | <4 | >0.99 |
| Sickle cell disease | 0 (0.00) | - | 0 (0.00) | - | 0 (0.00) | - | - | 0 (0.00) | - |
| Anemia (non-sickle cell) | <4 | - | <4 | - | <4 | - | - | <4 | - |
| Other | <4 | - | <4 | - | <4 | - | - | <4 | - |
| Reproductive disease | 7 (28.00) | 0 (0.00) | 7 (14.00) | **0.0003** | <4 | 5 (41.67) | 0 (0.00) | 7 (14.00) | 0.09 |
| PCOS | <4 | - | <4 | - | <4 | 0 (0.00) | - | <4 | - |
| Endometriosis | 0 (0.00) | - | 0 (0.00) | - | 0 (0.00) | 0 (0.00) | - | 0 (0.00) | - |
| Primary ovarian insufficiency | 0 (0.00) | - | 0 (0.00) | - | 0 (0.00) | 0 (0.00) | - | 0 (0.00) | - |
| Hysterectomy (full or partial) | 4 (57.14) | - | 4 (57.14) | **0.02** | 0 (0.00) | 4 (80.00) | - | 4 (57.14) | **0.0046** |
| Other | <4 | - | <4 | >0.99 | <4 | <4 | - | <4 | >0.99 |
| Endocrine/Metabolic disease | <4 | <4 | 4 (8.00) | 0.35 | <4 | <4 | <4 | 4 (8.00) | 0.42 |
| Thyroid | <4 | <4 | 4 (100.00) | 0.61 | <4 | <4 | <4 | 4 (100.00) | 0.68 |
| Smoking history | <4 | <4 | 5 (10.00) | >0.99 | <4 | <4 | <4 | 5 (10.00) | 0.84 |
| Current smoker? | <4 | <4 | <4 | >0.99 | 0 (0.00) | <4 | <4 | <4 | 0.58 |
| Past smoker? | <4 | <4 | <4 | >0.99 | <4 | 0 (0.00) | <4 | <4 | 0.58 |

### 2024–25 C.3 strains infect a younger population compared to C.5 subclade lineages but do not impact clinical symptoms or outcomes

To assess whether C.3.1 infections cause more severe disease than C.5 lineages, Demographic, co-morbidity, disease severity and disease outcome data was aggregated for IBV-infected patients presenting to clinics in the Johns Hopkins Hospital System in Baltimore, MD. 123 patients with complete influenza B genome sequences were considered **(Fig. S1)** and 10 genomes were excluded due to ≤98% coverage across all eight segments to ensure full IBV genomic coverage. This subset (n=113) was used to compare clinical and demographic characteristics between infections caused by C.3 (n = 67) and C.5.X subclades (C.5.1, C.5.6, and C.5.7; n = 46). The median age of patients infected with C.3 viruses was 8 years, significantly younger than those infected with C.5.X viruses (median 13 years) **(Fig. 1C-D)**. Notably, 40.3% of C.3-infected individuals were between 6–11 years of age, compared to only 17.4% in the C.5.X group **(Fig. 1C-D; Table S2)**. Disease severity and clinical outcome between C.3.1 and C.5.X were similar and beyond age, no demographic factors were significantly different infected individuals **(Table** S**2)**.

### 2024-25 C.3 strains encode a HA1:D197N introducing an additional N-linked glycosylation motif, and encode NA:E186R and NA:V395I mutations

HA segment sequence alignments revealed that all the 2024-25 strain belonging to the C.3 subclade contained an additional N-linked glycosylation (NLG) motif (N-X-S/T) due to a D197N substitution which is absent in the vaccine strain (B/Austria/1359417/2024) and circulating C.5.X subclades **(Fig. 1E -F).** In July, 2025 C.3 viruses encoding two reversion mutations, D197N and P208S, were assigned the C.3.1 clade^28^. Structural mapping of residue 197 onto the Influenza B virus hemagglutinin (PDB: 4FQM) revealed the site directly overlaps with the receptor binding site (RBS) and the 190-helix—critical regions for receptor attachment and neutralizing antibody recognition **(Fig. 1G**).

To assess whether the 2024–25 C.3 viruses acquire an additional N-linked glycan due to the HA1:D197N substitution, HA glycosylation was analyzed by western blot with and without PNGaseF treatment. In untreated samples, the C.3.1/re HA exhibited a higher molecular weight (∼80 kDa) compared with C.3 and C.5.1 HAs, consistent with the presence of an additional glycan **(Fig. S2)**. Following PNGaseF digestion, which removes N-linked glycans, all HA bands collapsed to a similar lower molecular weight (∼60 kDa), confirming that the observed size shift was attributable to differential N-linked glycosylation rather than amino acid substitutions alone **(Fig. S2)**. These results verify that C.3.1/re viruses encode an additional N-linked glycan at HA1:N197, introduced by the D197N mutation.

All C.3 genomes encode a V395I and a K186R substitution (**Fig. 1H**) in the NA gene that only appear together sporadically in IBV sequences until late 2024/early 2025 (**Fig. 1I**). The V395I mutation is located on the solvent-exposed 380-helix—a known antigenic site accessible to antibodies while K186R is proximal to the NA enzymatic active site and has been previously associated with reduced oseltamivir susceptibility when present in combination with 262T (**Fig.** 1J)^29,30^.

### C.3.1 subclade viruses in 2024-25 acquired 4 gene segments from C.5 lineages deemed C.3.1/re

Time-resolved maximum likelihood phylogenies were generated using complete genome sequences from the JHHS in combination with all available complete IBV genomes deposited to GISAID. To obtain finer resolution of parental ancestry among gene segments, the dataset was enriched during down sampling to include all C.3, C.3.1, and C.3.2 clade viruses deposited to GISAID between October 2020 and March 2026.

Concatenated genome phylogenies are sensitive visual approaches to reassortment detection through feature and topological incongruence, as segments inherited from disparate parental lineages can appear as long, or conflicting branching patterns compared to gene segments^31^. Concatenated genome phylogenies annotated by HA clade designation revealed that 2024–25 C.3.1 viruses originating from JHHS (n = 68) formed a highly divergent, monophyletic clade with a distinct ancestry from previously circulating parental C.3 viruses **(Fig. 2A).** The C.3.1 lineage was not directly connected to the parental C.3 clade in the phylogeny, suggesting a non-clock-like evolutionary rate or an unsampled intermediate ancestor.

**Figure 2.**
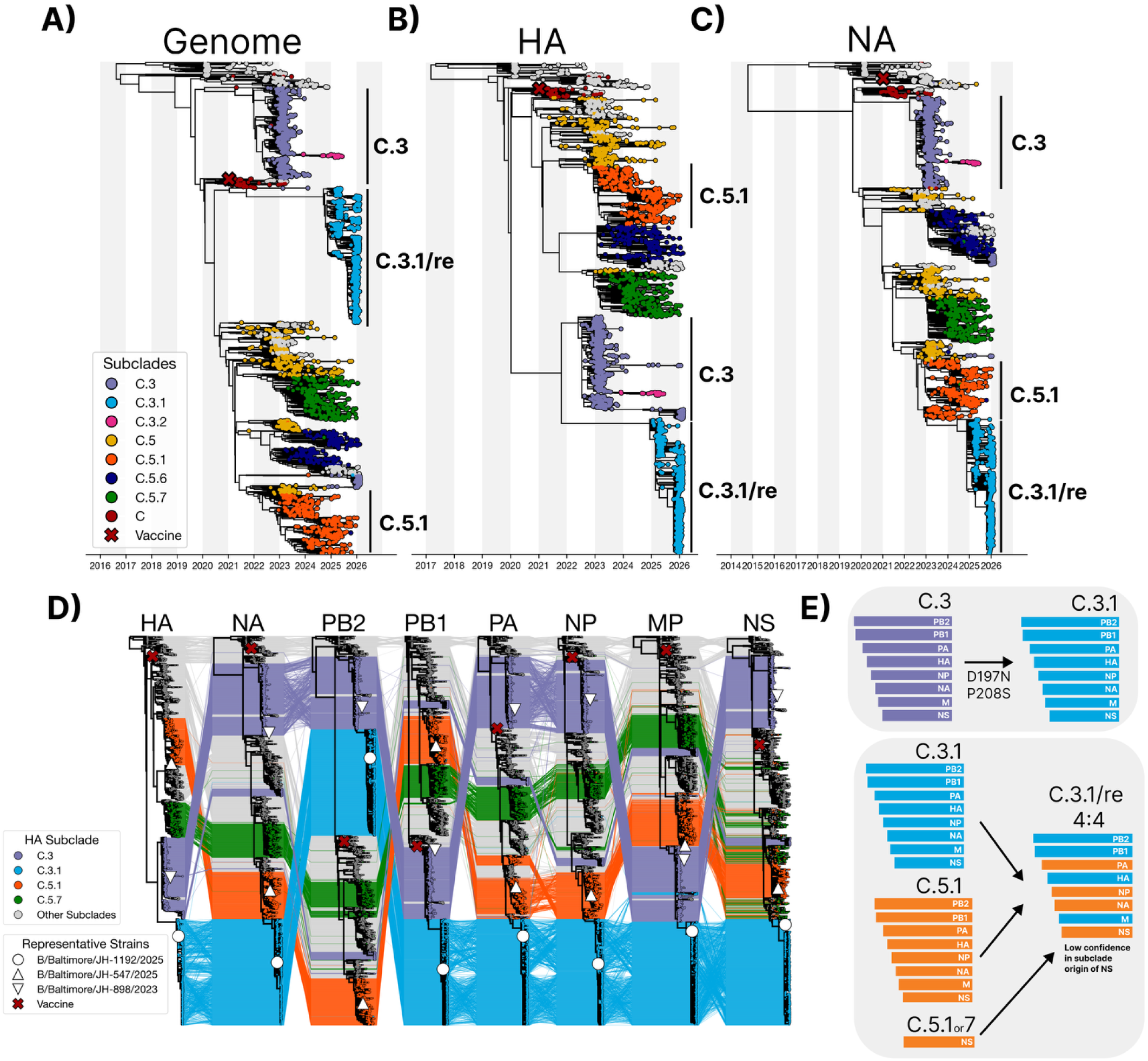
–2024-25 C.3 subclade viruses are 4:4 reassortment progeny with NA and internal segments derived from C.5.1 defined as C.3.1/re. **(A) C.3-enriched** maximum-likelihood phylogeny of concatenated genomes or HA **(B)** or NA **(C)** gene trees with tips colored by subclade. Major subclades (C.3, C.3.1, C.3.2, C.5, C.5.1, C.5.6, C.5.7, and C) are annotated by color and the 2024-25 vaccine strain tip (B/Austria/1359417/2021) is annotated by shape (X). Branch lengths were refined to inferred coalescent events using collection (reference strains) or sequencing date (JHH isolates) using augur refine. Tree tips belonging to C.3 in the concatenated **(A)** or NA **(C)** gene trees belong to separate reassortment events discussed in Figure 5. **(D)** A tanglegram consisting of 8 gene trees for each IBV segment interleaved and colored by HA subclade identity reveal a 4:4 reassortment with NA, PA, and NP clustering with the C.5.1 clade of each gene tree. The C.3.1/re NS segment clusters in a diverged cluster containing strains with definitive C.5.1 and C.5.7 HA membership. Auspice JSONs were ingested and visualized using Baltic. **(E)** A cartoon schematic representing the C.3 to C.3.1 clade advance and inferred segment ancestry of C.3.1/re from each of the 8 gene trees in relation to the clinical specimen’s HA subclade membership.

To determine the putative ancestry of individual genome segments, gene trees were constructed for all eight segments. In the HA phylogeny, 2024–25 C.3.1/re reassortant viruses formed a monophyletic clade directly linked to historical C.3 viruses, consistent with HA segment inheritance from C.3 **(Fig. 2B).** In contrast, the NA phylogeny revealed that these same C.3.1/re viruses clustered exclusively with C.5.1 sequences **(Fig. 2C),** indicating that all C.3.1/re viruses acquired their NA segment from a C.5.1 lineage.

To assess C.3.1/re ancestry of the internal segments, tanglegrams were constructed where the tree tips were colored by HA subclade identity and connected by lines to their position in each of the HA-NA-PB2-PB1-PA-NP-MP-NS phylogenies **(Fig. 2D).** Topological incongruence was identified in NA, PA, NP and NS segments where C.3.1 viruses shared direct ancestry with the C.5.1 lineage **(Fig. 2D and Fig. S3).** The precise ancestral clade determination for the NS segment is less clear due to limited genetic differences across C.5, C.5.7 and C.5.1 **(Figs S3-S4)**. A graphical summary of C.3.1/re genome composition is shown in **Fig. 2E**, indicating C.3.1/re as a 4:4 reassortant virus with 4 segments originating from a C.3.1 ancestor (PB2, PB1, HA, M) and 4 segments likely originating from a C.5.1 ancestor (PA, NP, NA, and NS

### 2024-25 C.3.1/re strains are antigenically drifted from circulating C.5.1 and the 2024-25 vaccine strain

To assess where baseline or post vaccination serum antibody titers recognize C.3.1/re, serum neutralization assays using clinically isolated infectious viruses were performed against the 2024–25 vaccine strain (B/Austria/1359417/2024; subclade C), a parental C.3 virus from the 2022-23 season (B/Baltimore/JH-898/2023), a C.5.1 virus from the 2023–24 season (B/Baltimore/JH-547/2024), and a representative C.3.1/re virus from 2024–25 (B/Baltimore/JH-1192/2025) at the time of vaccination (day 0) and 28 days post-vaccination ( Table S3-S4). A cohort of 50 individuals from the 2024-25 Johns Hopkins Center of Excellence for Influenza Research and Response (JH-CEIRR) Vaccine Cohort (Table 2) was used to assess population immunity. Baseline (Day 0) neutralizing antibody titers quantified by NT50 were similar amongst the homologous vaccine, C.3, and C.5.1 strains (Fig. 3A). However, NT50 titers against C.3.1/re were significantly lower compared to the vaccine and parentals with 84% of individuals (42/50) having no detectable neutralizing antibodies. Following vaccination, NT50 titers were markedly lower to C.3.1 with 78% of individuals (38/50) with geometric mean titers below the limit of detection (Fig. 3B and Table 2). Seroconversion rates (≥4-fold NT50 increase) to the vaccine,C.3 and C.5.1 parental strains were similar at 38%, 42%, 36%, respectively. However, only one individual (2%) seroconverted to C.3.1/re following vaccination (Fig. 3C). An area under the curve analysis was used to assess fold change between baseline and vaccine geometric mean titers. The mean increase in neutralization AUC titer after vaccination rose 1.2 fold for C.3.1/re (Fig. 3D) while titers increased nearly 8-fold for the vaccine and 6.2-fold for the C.5.1 strain and 7.7 for the C.3 strain. C.3.1/re AUC titers were significantly different by birthyear with the highest titers in those born between 1960 and 1970 (Fig. 3E and Fig. S5). These results indicate the C.3.1 /re viruses can evade preexisting population immunity as well as vaccine induced immunity.

### Removal of the N-linked glycosylation motif at HA residue 197 restores neutralizing activity of vaccine sera to C.3.1/re

One of the C.3.1/re clade defining mutations, D197N, is considered a potential contributor to antigenic drift as it may shield a wide footprint of 190-helix neutralizing antibodies. To investigate this, we generated an infectious clone of C.3.1/re mutating the asparagine at residue 197 to the parental aspartic acid thus removing the putative NLG motif spanning residues 197-199 (NET to DET). This virus is referred to as rg-C.3.1/re:N197D where rg = reverse genetics. Using the same vaccination sera to assess clinical isolates, we observed restoration of neutralizing antibody titers to rg-C.3.1/re:N197D with NT50 GMT values comparable to the Vaccine, C.3 and C.5.1 (Fig. 3A-3B). Baseline and post vaccination titers were observed to be significantly higher than the C.3.1/re clinical isolate. Furthermore, seroconversion rates and AUC fold-change were significantly boosted to those like C.5.1, 36% and x6.5, respectively (Fig. 3C-D). This suggests that in our cohort, neutralizing antibodies both at baseline and post vaccination with B/Austria/13594172021, were focused near or around the site 197 (190 helix). When baseline and postvaccination titers were analyzed with respect to birth year, there was a significant association of older age with neutralization of C.3.1/re, indicating that individuals born after 1970 were more likely to lack neutralizing activity (Fig. 3E).

### C.3.1 /re viruses encode a C.5.1 NA with lower sialidase activity compared to the parental C.5.1 but remain sensitive to oseltamivir

To determine whether the C.3.1/re NA acquired via reassortment with a C.5.1 parent has altered sialidase activity, NA activity and oseltamivir sensitivity was compared among representative C.3.1/re and C.5.1 isolates (**Fig. 3F-G**). The C.3.1/re virus displayed reduced total NA activity per unit of infectious virus compared to its C.5.1 parent (**Fig. 3F**). Relative reduction in luciferase activity was quantified by normalizing luciferase readouts such that no oseltamivir (i.e., infectious virus alone) represented 0% inhibition of NA activity (NAI), and no virus (i.e., assay buffer alone) represented 100% inhibition of NA activity. Representative C.3.1 /re and C.5.1 viruses showed similar IC_50_, IC_80_ and IC_90_ concentrations. These data show that while the C.3.1/re NA has slightly reduced sialidase activity relative to parental C.5.1 NA (**Fig. 3F**), both viruses show comparable sensitivity to oseltamivir (**Fig. 3G**).

**Figure 3.**
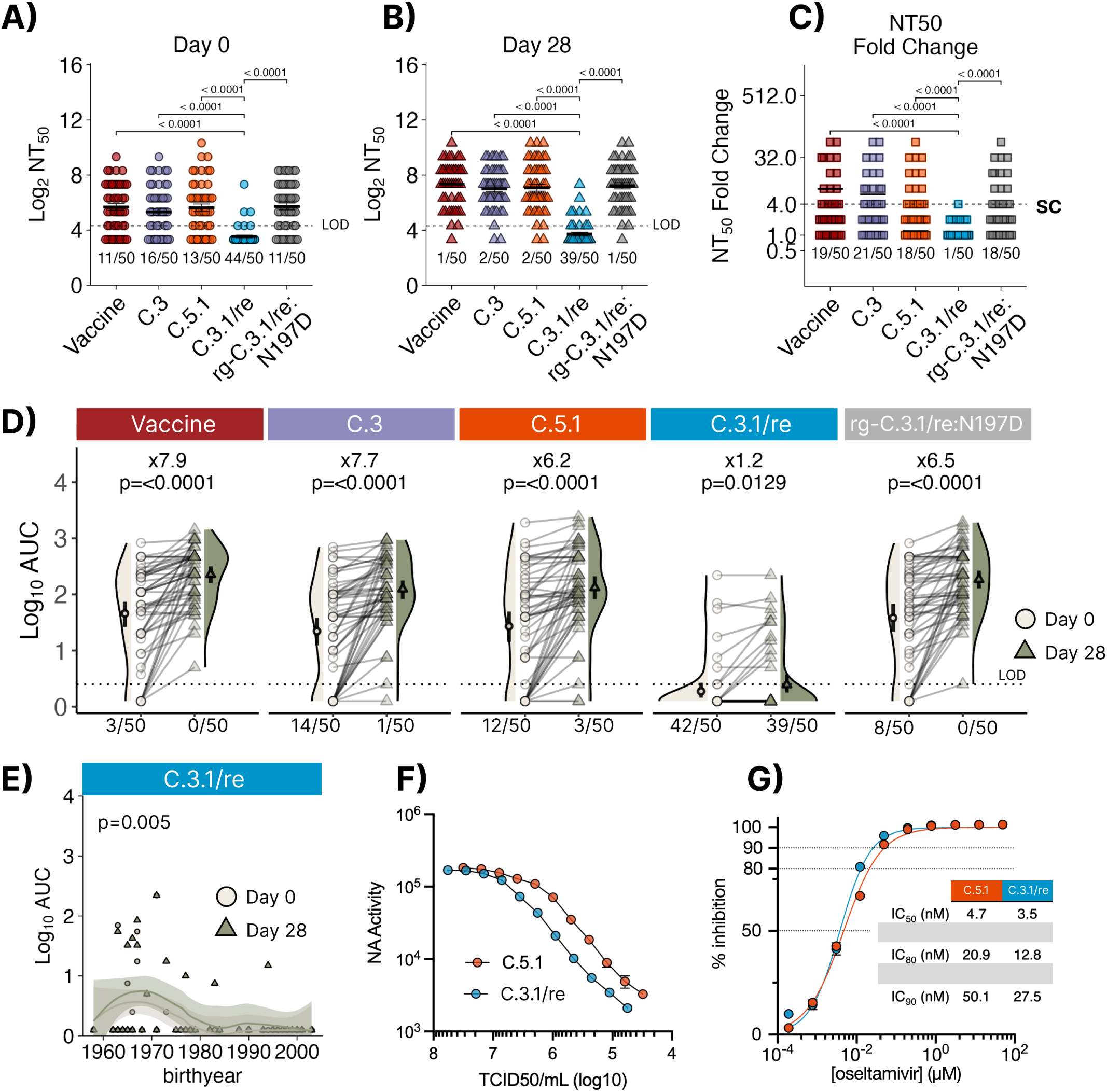
The C.3re subclade is antigenically drifted compared to parental viruses and encodes a neuraminidase with lower sialidase activity per infectious unit. The Johns Hopkins CEIRR Network influenza vaccine cohort consisted of 50 individuals receiving the 2024-25 trivalent Northern Hemisphere influenza vaccine formulation. Serum neutralizing antibody titers (NT50 ) with mean and standard error plotted **(A)** at the time of vaccination (Day 0) and **(B)** 28 days post vaccination against the 2024-25 B/Victoria vaccine strain (B/Austria/1359417/2021 subclade C) and a representative 2024-25 C.5.1 dominant parental (B/Baltimore/JH-547/2024 subclade C.5.1) or a representative 2024-25 C.3.1/re reassortment virus (B/Baltimore/JH-1192/2025). Each point on the graph represents an individuals’ observed titer where circles denote Day 0 and triangles denote Day 28. The number of seronegative individuals over the total number of individuals analyzed are shown under the datapoints for each virus. **(C)** NT50 fold change for day 0 to day 28 was used to determine seroconversion which was defined as a NT50 ≥ 4. Individuals who seroconverted are shown as a fraction out of 50 beneath each data group. The dotted line represents the limit of detection (LOD), defined as the lowest value considered to be NT50 positive at the starting dilution of 1/20. No neutralizing activity is graphed as one-half the LOD. **(D)** Area under the curve (AUC) was calculated and used to show changes in titers at day 0 and day 28 post vaccination with an LOD set to 2.5. AUC fold change is shown above pairings. The lines connect the same individual’s AUC titers between collection timepoints. The number of seronegative individuals over the total number of individuals analyzed are shown under the datapoints for each virus. Statistical analysis within or between non-transformed NT50 or AUC timepoint values was performed using paired Wilcoxon tests with Bonferroni correction post-hoc. Adjusted p-values are shown above significantly different comparisons. **(E)** Log10 AUC plotted against birthyear for C.3.1/re. Each point represents a single participant AUC titer at baseline or 28-days post vaccination. Timepoint AUC values were fit against a non-linear model (loess). Kruskal-Wallis nonparametric test with Bonferroni’s correction for age analyses. **F)** Neuraminidase (NA) activity was measured via NA-Star assay across serial dilutions of viral seed stocks. Oseltamivir resistance **(G)** was evaluated using the NA-Star assay. The amount of virus used in each reaction corresponded to the dilution that produced approximately 10⁴ RLU. Each symbol represents the mean of eight biological replicates, and error bars denote standard deviation. Inhibitory concentrations (IC₅₀, IC₈₀, and IC₉₀) were derived from nonlinear regression curves fitted using a log(agonist) vs. normalized response model.

### C.3.1 /re persists in the United States with few detectable cases globally

To generalize relative C.3.1/re subclade abundance amongst global regions, all available IBV HA sequences from GISAID between October 2020 and March 18, 2026 were assigned HA clade designations and the relative proportions of all subclades were quantified by global region: North America, South America, West Asia, South Asia, East Asia, Korea/Kapan, Africa, Europe and Oceania **(Fig. 4)**.

**Figure 4.**
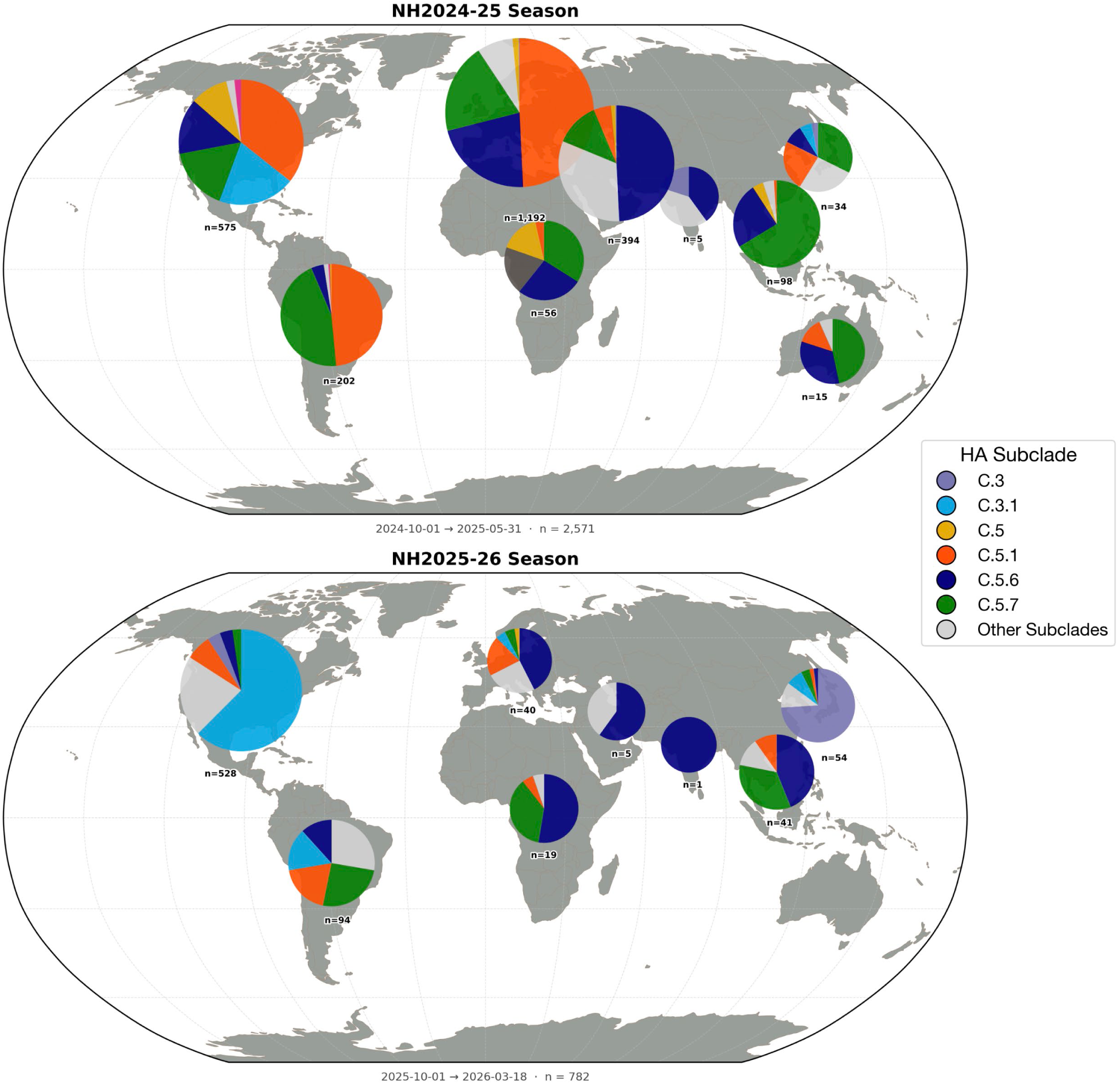
Global distribution of Influenza B subclades across two subsequent Northern Hemisphere Influenza seasons as of March 18, 2026. Bubble size represents relative sequence abundance of IBV hemagglutinin sequences deposited to GISAID between October 2020 and March 18, 2026. Colors represent the fraction of HA subclade abundance as assigned by Nextclade (see methods). The major circulating C.5 lineages and C.3 lineages are highlighted. Other minor circulating clades are colored grey with more detailed regional subclade abundance over time outlined in Supplemental Figure 1. Bubble placement is defined by region: North America, South America, Europe, Africa, West Asia, East Asia South Asia, Japan/Korea and Oceania.

Throughout the late 2024-25 season, C.3.1/re was only detected in the North America (**Fig. 4**), primarily in the eastern United States. Towards the end of the 2024-25 season, C.3.1/re genomes originating from Japan (representative B/TOKYO/EIS-13-175/2025) were reported **(Fig. 4; Fig. S5)**. In the 2025-26 season, C.3.1 displaced C.5.1 in North America and has been detected in South America, Europe, Japan and Korea. Most C.3.1/re sequences originate from North America while C.3.1/re appears to be a minor subclade globally As of March 18, 2026, global genomic surveillance data indicate that C.3.1/re viruses were most prevalent in the United States, where they represent the majority of recent IBV detections (**Fig. 3**). Although all IBV subclades appear globally in varying magnitude, the C.5.1 subclade is disproportionately represented in Western regions, whereas the C.5.6 and C.5.7 lineages are more prevalent in Eastern regions **(Fig. 4)**.

### Additional C.3 reassortments with C.5 lineages is concurrent with a HA:D197N reversion

As of March 18, 2026, the IBV C.3 lineage consists of the following clade designations: C.3, C.3.1 and C.3.2. Following the emergence of the C.3.1/re reassortment event, 2 additional reassortment events where a C.3 HA segment is retained in a C.5-lineage backgrounds (**Fig. 5A-B).** The first noted event, classified here as C.3/re, encodes a C.3 HA with D197N and shares direct ancestry with C.5.6 while encoding internal segments from either C.5 or C.5.7 parental viruses (**Fig. 5A-B**). The C.3/re reassortment event was first detected in Japan in the 2025-26 season and has continued to increase in detection frequency. Putative ancestry for non-HA segments was resolved by constructing HA tanglegrams connecting taxa in the corresponding NA-PB2-PB1-PA-NP-MP-NS phylogenies **(Fig. 5B).** The ancestry of non-HA segments in C3/re maps to a putative (C.5.6)-(C.5.6)-(C.5)-(C.5)-(C.5)-(C.5.7)-(unknown), (NA-PB2-PB1-PA-NP-MP-NS), constellation where the NS segment clusters tightly with a clade populated with C.5, C.5.1, and C.5.7 HA-assigned taxa **(Fig. 5B).** A second and minor reassortment event, named C.3/re.1, was detected in the United States in the 2025-26 season (**Fig. 5A and B**). C.3/re.1 encodes a C.3 HA with the D197N and is a putative C.3 and C.5 reassortment with a genome constellation of (C.5)-(C.5)-(C.5)-(C.5)-(C.3)-(C.5)-(C.3), (NA-PB2-PB1-PA-NP-MP-NS).Both reassortment events donate a C.3 HA segment encoding a D197N and therefore a NLG motif.

**Figure 5.**
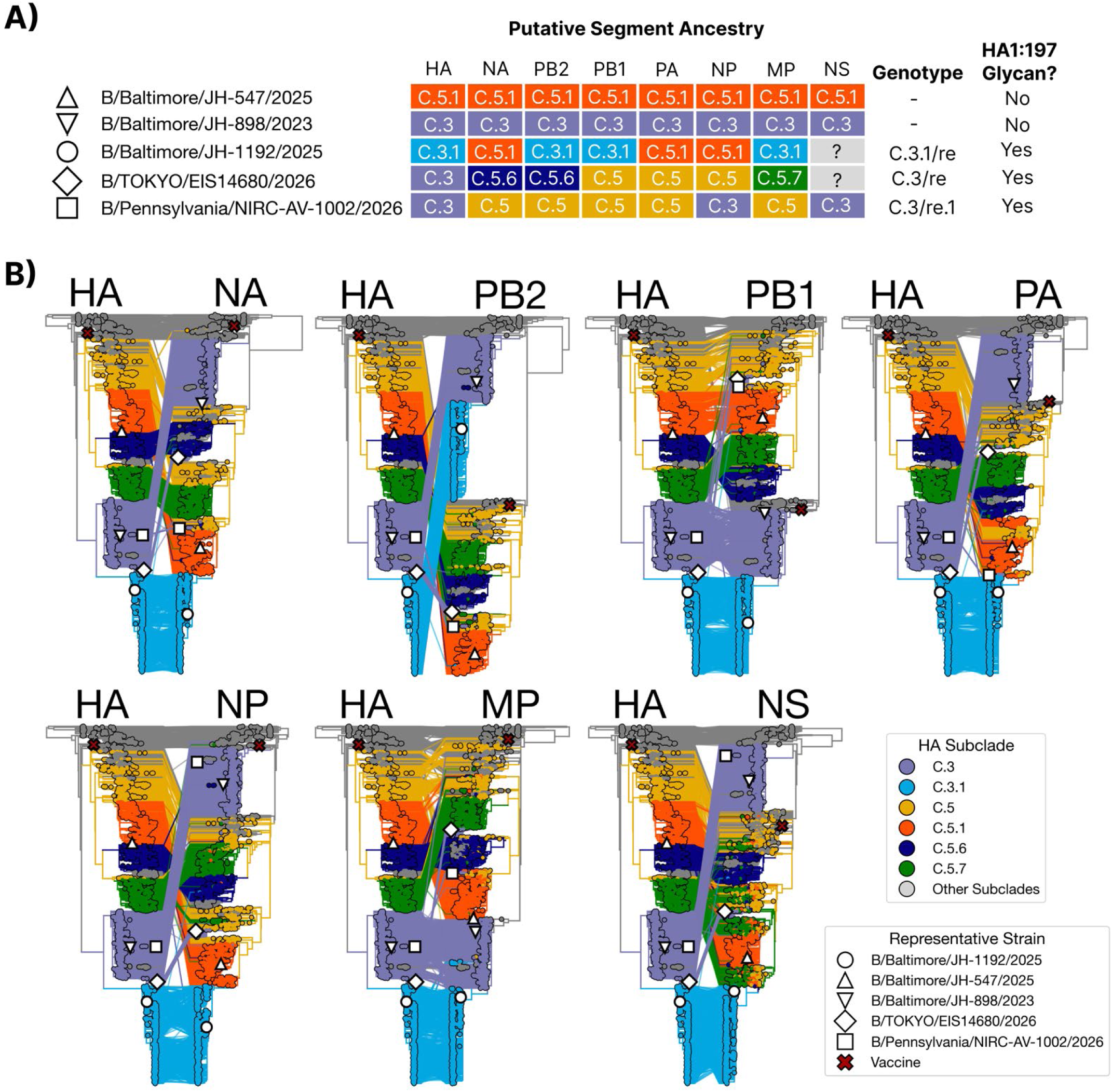
Putative ancestries of 2 additional reassortment events denoted at C.3/re and C.3/re.1. **(A)** Genome-identity schematics of the non-reassorted C.5.1 or C.3 parental genomes and corresponding C.3.1/re, C.3/re and C.3/re.1 reassortment segment constellations with their corresponding genotype name if reassorted. Each parental virus or reassortment event is a corresponding shape and representative strain which corresponds to its location in each of the 8 segment phylogenies **(B)**. Gene-level phylogenies of all 8 segments were constructed and compared pairwise with the HA gene tree. Segments were assigned HA subclade ancestry based on the HA subclade assignment of their most recent common ancestor (MRCA) within each segment tree. We assigned tree tip colors by HA subclade identity. HA subclade identity was transferred from the HA tree to the same tip in each of the 7 remaining segment trees tree. Tree tips are colored by HA subclade identity and connected by lines corresponding to their placement in the annotated gene phylogeny. Phylogenies were visualized in Baltic and formatted in Figma 125.9.10

## DISCUSSION

Genomic surveillance of seasonal influenza viruses remains essential for epidemic preparedness and vaccine strain selection. During the 2024–25 influenza season, the C.3.1 subclade displaced the previously dominant C.5.1 lineage in Baltimore, MD. Notably, the reassorted C.3.1/re virus disproportionately infected younger individuals (median age 8 years), with 40.3% of cases occurring in children aged 6–11 years. This age distribution is consistent with the broader epidemiology of B/Victoria viruses, which are known to preferentially infect pediatric populations, but such skewing has not been well described between B/Victoria subclades ^32,33^. The known epidemiological tendency of pediatric infection combined with an antigenically naïve population may explain the differences in age see between C.3.1/re and C.5.X in the 2024-25 JHHS season. However, our neutralizing antibody data with an adult population would indicate a broad C.3.1/re susceptibility across age groups. Despite likely prior exposure to B/Victoria viruses, most adults in our cohort had no detectable neutralizing antibody titers against C.3.1/re either before or after vaccination. This indicates that C.3.1/re effectively escaped both preexisting and vaccine-induced immunity, even in an ostensibly experienced adult population. One contributing factor may be immune imprinting toward epitopes present in recently circulating or vaccine-matched strains, particularly those lacking glycosylation at HA site 197. The absence of serum neutralization against C.3.1/re, contrasted with preserved responses to C.5.1, supports the idea that antigenic drift—rather than age alone—underlies the observed susceptibility patterns. The glycosylation at HA position 197 in C.3.1/re likely masks immunodominant epitopes that were exposed in B/Austria/2021-like viruses, thereby undermining both infection- and vaccine-induced immunity. Further investigation as to the potential immuno-focusing consequences of repeated vaccination and exposure to IBVs which lack the 197 NLG is warranted ^34^.

The addition of an N-linked glycan in the hemagglutinin (HA) receptor binding site (RBS) is a well-known determinant of epitope masking in Influenza A and B viruses ^5,21^. Antigenic changes driven by a gain or loss of glycosylation at IBV HA reside 197 have been documented previously and are a known egg adaptation mechanism for IBV vaccine strains ^26^. Historically, B/Victoria lineage viruses retained glycosylation at this site until the emergence of B/Austria/2021-like viruses, which lost this modification ^34–39^ . The loss of glycan resulting from egg-adaption of IBV vaccine strains leads to the production of antibodies with poor recognition of circulating, glycosylated IBV strains ^27,40^. Consistent with prior studies, we find that the presence of this glycan substantially reduces neutralizing antibody recognition, while its removal restores neutralizing antibody titers.

Intra-subtype reassortment between human influenza viruses has been previously observed to increase disease severity while further diversifying the repertoire of influenza gene segment combinations ^31,41^. Phylogenetic analyses revealed that C.3.1/re viruses acquired multiple internal gene segments from co-circulating C.5.1 viruses, including the C.5.1 NA segment which carries a surface-facing V395I mutation within the 380 helix antigenic site that may confer escape from some NA binding antibodies ^42^. The significance of reassortment on IBV fitness is clear, but not explored well mechanistically ^17,18^.

IBV is known to have slow global migration with epidemic models supporting differences in age of infection as a likely driver of regional persistence and differences in patterns of global circulation ^4,19^. To our knowledge, the C.3.1/re subclade—corresponding to the C.3.1 HA designation—was first detected in North America. Our analyses do not indicate that C.3.1/re viruses have gained a competitive advantage over C.5 lineages in the Southern Hemisphere; C.3.1/re detections remain rare or absent in available surveillance data. However, the gain of an HA 197 glycosylation site and intraclade reassortment has occurred independently at least three times, suggesting some mechanistic advantage of both to IBV fitness and circulation.

The emergence of C.3.1/re occurred after IBV vaccine strain selections were made for the 2025-26 Northern Hemisphere influenza vaccine and therefore, it was not available to be considered as a vaccine strain candidate ^37^. This highlights the need to shorten the time between influenza vaccine strain selection and the initiation of the fall seasonal influenza vaccine campaign to fully identify and characterize late season virus variants and allow them to be considered as vaccine candidates. The implementation of mRNA vaccine platforms could enable rapid updates against mismatched strains detected after the annual February and September WHO vaccine selection meetings. This approach could improve strain concordance and reduce infections, hospitalization and deaths ^43^.

The 2024–25 Northern Hemisphere IBV vaccine component was subclade C B/Austria/1359417/2024 (V1A.3a.2), used since 2022. Interim VE, largely against early-season C.5.X viruses, was 58% in Europe but could not be estimated in the U.S. due to limited cases ^44,45^. IBV comprised 17.2% of positives (week ending in Jun 28, 2025) ^46^. Egg-adapted ferret antisera recognized 96.6% of isolates by HI, mainly C.5.X ^47^. Our results would predict that overall, VE against IBV will drop towards the end of the 2024-25 influenza season with the emergence of C.3.1/re with a similar drop in the 2025-26 season. For the upcoming 2026-27 Northern Hemisphere season, the selected IBV vaccine egg component lacks a putative NLG as reported by the WHO (B/Tokyo/EIS13-175/2025-E3/SpE1; EPI_ISL_20373028) which is likely to induce neutralizing antibodies that are blocked by C.3.1 HA containing viruses that have the D197N NLG.

This study has several limitations. First, our genomic dataset reflects sequencing efforts in a single hospital system and may not fully capture community transmission dynamics. Broader population-based surveillance was limited, and global representation in publicly available databases such as GISAID remains uneven, potentially obscuring the true geographic distribution of C.3.1/re viruses. Not all IBV positive cases underwent whole-genome sequencing, and thus clade assignments were available for only a subset of patients. Furthermore, time-scaled phylogenies relied on maximum likelihood methods rather than Bayesian inference, which may underestimate uncertainty in time to the most recent common ancestor (tMRCA) estimates and geographic movement.

## MATERIALS AND METHODS

### Ethical considerations and human subject approval

The Johns Hopkins Institutional Review Board has approved this work. The research was performed under protocols IRB00091667 and IRB00331396. Genomes are available in the Global Initiative on Sharing All Influenza Data (GISAID) database. Accession numbers available in **DOI XXX**. Serologic samples for this study were obtained from healthcare workers (HCWs) recruited from the Johns Hopkins Centers for Influenza Research and Response (JH-CEIRR) during the annual Johns Hopkins Health System employee influenza vaccination campaign in the Fall of 2024. Serum was collected from subjects at the time of vaccination and approximately 28 days later. Virus was isolated for this study from deidentified IBV positive nasal swabs under the JHU School of Medicine Institutional Review Board approved protocol, IRB00288258.

### Acute infection study population

Standard-of-care diagnostic influenza testing was conducted at the JHHS. Detection of influenza virus was performed with either the Cepheid Xpert Xpress SARS-CoV-2/Flu/respiratory syncytial virus test (Sunnyvale, CA, USA) or the ePlex RP/RP2 respiratory panels (Roche Diagnostics, Indianapolis, IN, USA). The Xpert Xpress assay targets the matrix the M and NS segments for IBV. Clinical samples were collected between December 2024 and April 2025, and corresponding clinical and demographic metadata were extracted in bulk through JHHS electronic medical charts.

### Nucleic Acid Extraction and Whole Genome Amplification from clinical specimens

Nucleic acid was extracted using the Chemagic Viral RNA/DNA Kit following the manufacturer’s instructions (Revvity, Waltham, MA, USA). The whole genomes of IBV were amplified using Invitrogen Supercript III (Waltham, MA, USA) and universal IBV primer cocktail.^49^ Library preparation was performed with NEBNext Artic SARS-CoV-2 Companion Kit (New England Biolabs, Ipswich, MA, USA), and sequencing was performed following manufacturer’s instructions, using R10.4.1 flow cells on a GridION (Oxford Nanopore Technologies, Oxford, UK).

### Influenza Genome Assembly

Fastq files were demultiplexed using the artic_guppyplex tool (Artic version 1.2.2). Nucleotide sequence assembly was performed using the default settings of the FLU module of the Iterative Refinement Meta-Assembler (IRMA version 1.0.2) which include a minimum average quality score of 24 and a site depth of 100. The alignment of genomes and reference sequences, downloaded from GISAID, was performed using the built-in alignment tool in Nextclade. Quality control scores for sequences were assigned using the built-in pipeline available in Nextclade. Sequences with overall quality scores of 30 and above were excluded from sequence analysis.

### Cell Culture and media

Madin-Darby canine kidney (MDCK) cells (provided by Dr. Robert A. Lamb, Northwestern University), MDCK-SIAT-1 cells (provided by Dr. Scott Hensley, University of Pennsylvania) and MDCK-SIAT-1-TMPRSSII cells (provided by Dr. Jesse Bloom, Fred Hutchinson Cancer Center) were maintained in complete medium (CM) consisting of Dulbecco’s Modified Eagle Medium (DMEM) supplemented with 10% fetal bovine serum, 100 units/mL penicillin/streptomycin (Life Technologies) and 2 mM Glutamax (Gibco) at 37 °C and 5% CO2. Human nasal epithelial cells (hNEC) (PromoCell) were cultivated as previously described ^6^. Infection media consisted of 0.3% bovine serum albumin,100 units/mL penicillin/streptomycin (Life Technologies), 2 mM Glutamax (Gibco) and 2.5 µg.mL of N-acetylated trypsin (NAT) from bovine pancreas (Sigma) for MDCK-SIAT-1 infections only (not MDCK-SAIT1-TMPRSS2 or Human Nasal Epithelial cell infections).

### Virus Isolation on Human Nasal Epithelial cells (hNECs) and MDCK-SIAT-1-TMPRSSII cells

Nasopharyngeal swabs or nasal washes from influenza B–positive individuals were used for virus isolation on MDCK-SIAT-1-TMPRSSII cells or primary human nasal epithelial cell (hNEC) cultures as previously described ^6,10^. MDCK-SIAT-1-TMPRSSII cells were maintained CM. For isolation, specimens were inoculated onto cells, washed, and incubated at 33°C in IM lacking NAT. Supernatants were harvested, and infectious titers were quantified by TCID50 and were expanded to generate seed stocks.

### Virus plasmid cloning and recombinant virus generation of eg-C.3.1/re:N197D

An eight-plasmid reverse genetics system (pDP2002)^50,51^ was used to rescue recombinant viruses corresponding to B/Baltimore/JH-1192/2025 (C.3.1/re). To evaluate the contribution of HA residue 197 glycosylation to antigenicity, the HA plasmid was site-directed mutagenized to convert N197 to D (NET→DET), thereby removing the N-linked glycosylation motif at residues 197–199. All plasmids and rescued viruses were sequence-verified across the coding regions of the modified segment(s). Detailed procedures for cloning and site directed mutagenesis are in the supplemental methods.

### Virus Stock Preparation

MDCK, MDCK-SIAT-1, and MDCK-SIAT-1-TMPRSSII cells were maintained in CM at 37 °C and 5% CO2. Primary human nasal epithelial cells (hNECs; PromoCell) were cultured as previously described ^6^. Influenza B viruses were isolated from IBV-positive nasopharyngeal swabs or nasal washes using MDCK-SIAT-1-TMPRSSII cells or hNEC cultures (see supplemental methods). Infectious titers were quantified by TCID50 on MDCK-SIAT-1 cells (Reed–Muench).

### Tissue Culture Infectious Dose 50 (TCID50) Assay

MDCK-SIAT-1 cells were seeded in a 96-well plate 2 days before assay and grown to 100% confluence. Cells were washed twice with PBS+ then 180 µL of was added to each well. Ten-fold serial dilutions of virus from 10^−1^ to 10^−7^ were created and then 20 µL of the virus dilution was added to the MDCK-SIAT-1 cells. Cells were incubated for 6 days at 33 °C then fixed with 2% formaldehyde. After fixing, cells were stained with Naphthol Blue Black, washed and virus titer was calculated using the Reed and Muench method.

### Validating hemagglutinin N-linked glycosylation motifs by PNGAseF treatment and western blot

To confirm glycosylation differences associated with HA residue 197, MDCK-SIAT-1 cells were infected (MOI 1) and lysates were analyzed by SDS-PAGE and immunoblot with an anti–influenza B HA monoclonal antibody Thermo Fisher MA5-29901 (see supplemental methods). Paired lysates were treated ± PNGase F prior to electrophoresis to distinguish glycosylation-dependent mobility shifts.

### Serum Neutralization Assay (NT_50_)

Day 0 and day 28 serum samples from JH-CEIRR participants were treated with receptor-destroying enzyme (Denka-Seiken), heat inactivated and serially diluted two-fold. Diluted sera were incubated with 100 TCID50 of infectious virus and then used to infect confluent MDCK-SIAT-1 cells. Following a 1 hour, incubation serum+virus was replaced with IM containing N-acetyl trypsin (2.5 µg/mL) and incubated for 6 days before they were fixed and stained as previously described ^14^. Neutralization titers (NT50) were defined as the highest serum dilution associated with ≥50% reduction in cytopathic effect.

### Assessment of NA activity and oseltamivir sensitivity

Relative neuraminidase (NA) activity and oseltamivir susceptibility were assessed using the NA-Star–based Influenza Neuraminidase Inhibitor Resistance Detection Kit (Thermo Fisher) according to the manufacturer’s protocol. Virus input was normalized by NA-Star per infectious unit (TCID50). Oseltamivir carboxylate (MedChemExpress) was serially diluted and incubated with virus prior to NA-Star substrate addition. Percent NA inhibition was calculated by normalizing luminescence to virus-only (0% inhibition) and no-virus (100% inhibition) controls. IC50 values were estimated by nonlinear regression in GraphPad Prism.

### Influenza B clade and genotype definitions

Influenza B viruses were classified using algorithmic clade proposals for the HA segment implemented into Nextclade (last accessed March 18, 2026).^52–54^ As of March 18, 2026, the recognized C.3 subclades and their clade defining amino acid mutations include C.3 (128K, 154E), C.3.1 (197N, 208S), and C.3.2 (197N, 208P) available at: https://github.com/influenza-clade-nomenclature/seasonal_B-Vic_HA/tree/main/subclades.

### Influenza B Reassortment Nomenclature

Reassortment was inferred by comparing phylogenies of all eight influenza B virus genome segments. Viruses were classified as reassortants when segment trees were incongruent, specifically when the HA clustered within the C.3/C.3.1 clade while one or more non-HA segments clustered within the C.5 lineage. Reassortant genotypes were labeled using the format “<HA clade>/re[.X]”, where “/re” denotes reassortment and an optional numeric suffix distinguishes independent reassortment events within the same HA clade; designations were updated as HA clade definitions were refined. Complete rationale is available in the supplemental methods.

### Protein Structure modeling

Mutations in Influenza B **Hemagglutinin** (PDB 4FQM) and Neuraminidase (PDB 4CPL) were visualized using Protein Imager^55^ .

### Phylogenetic Reconstruction of C.3.1/re ancestry using Maximum Likelihood

19,689 Influenza B genomes were access from GISAID filtered by original passage and collection date between October 2020 and August 22, 2025. 3836 whole genomes were down sampled by collection week and country implement in augur filter (see <u>01_ingest.qmd</u>) ^56^. Concatenated genome- and gene-level phylogenetics (totaling to 9) were constructed using IQ-Tree2 available in the augur tree module ^23,24^ and resulting branches were refined using treetime^56,58^. **(Fig. S3**). Nextclade was used to call clade and subclade designations (last accessed March 18^th^, 2026) sequence quality metrics, and putative glycosylation sites^53^

### Statistical analyses

Statistical analyses were performed in R (4.3.2) using Tidyplots ^59^ or Graphpad Prism (10.4.2) and described in figure legends Time to the most recent common ancestor estimates and confidence intervals were performed using baltic^17^.

## Data availability

Genomes queried from the Global Initiative on Sharing All Influenza Data used in this study are available at https://doi.org/10.55876/gis8.260402we (EPI_SET_260402we). Genomes generated in from the Johns Hopkins Hospital Network used in this study are available at: https://doi.org/10.55876/gis8.260511na (EPI_SET_260511na)

All scripts generated in this publication are available athttps://github.com/elginakin/influenzab_c.3. Interactive gene and genome-level phylogenies Nine genome- and gene-level phylogenies are publicly accessible through Nextstrain Groups at https://nextstrain.org/groups/PekoszLab-Public/akine/ibvc3/vic/genome. All datasets used and/or analyzed during the current study are available through the Johns Hopkins Research Data Repository at **[TODO: INSERT DOI WHEN RELEASED]**.

## Acknowledgments

This work was supported by the National Institutes of Health (NIH) contract 75N93021C00045 Johns Hopkins Centers of Excellence in Influenza Research and Response, and NIH T32 AI007417. The authors thank the healthcare workers who enrolled and participated in the study. We are grateful for the efforts of the clinical coordination team at the Johns Hopkins Hospitals who collected samples. We gratefully acknowledge all data contributors, i.e., the Authors and their Originating laboratories responsible for obtaining the specimens, and their Submitting laboratories for generating the genetic sequence and metadata and sharing via the GISAID Initiative, on which this research is based. A list of laboratories who contributed sequences for strains analyzed in this work is provided through GISAID **EPI_SET_260402we**. The 2024-25 Influenza B vaccine strain component, B/Austria/1359417/2024 was kindly provided by Dr. John Steel through the US Centers for Disease Control (CDC). EA’s participation as a student in the 2025 Workshop on Molecular Evolution (MOLE) at the Marine Biological Laboratory in Woods Hole, MA provided important insight into constructing phylogenies and reassortment networks pivotal to this study. We thank the laboratories of Heba Mostafa and Andrew Pekosz for discussion of data and future directions.

## References

1. van de Sandt CE, Bodewes R, Rimmelzwaan GF, de Vries RD. Influenza B viruses: not to be discounted. Future Microbiol 2015;10(9):1447–65.

2. Costa JC da, Siqueira MM, Brown D, et al. Vaccine Mismatches, Viral Circulation, and Clinical Severity Patterns of Influenza B Victoria and Yamagata Infections in Brazil over the Decade 2010-2020: A Statistical and Phylogeny-Trait Analyses. Viruses 2022;14(7):1477.

3. Dawood FS. Interim Estimates of 2019–20 Seasonal Influenza Vaccine Effectiveness — United States, February 2020. MMWR Morb Mortal Wkly Rep [Internet] 2020 [cited 2025 July 11];69. Available from: https://www.cdc.gov/mmwr/volumes/69/wr/mm6907a1.htm

4. Bedford T, Riley S, Barr IG, et al. Global circulation patterns of seasonal influenza viruses vary with antigenic drift. Nature 2015;523(7559):217–20.

5. Ni F, Kondrashkina E, Qinghua Wang, Qinghua Wang, Wang Q. Structural basis for the divergent evolution of influenza B virus hemagglutinin. Virology 2013;446(1):112–22.

6. Swanson NJ, Marinho P, Dziedzic A, et al. 2019–2020 H1N1 clade A5a.1 viruses have better in vitro fitness compared with the co-circulating A5a.2 clade. Sci Rep 2023;13(1):10223.

7. Wang Y-F, Chang C-F, Chi C-Y, Wang H-C, Wang J-R, Su I-J. Characterization of glycan binding specificities of influenza B viruses with correlation with hemagglutinin genotypes and clinical features. J Med Virol 2012;84(4):679–85.

8. Nakagawa N, Kubota R, Maeda A, Okuno Y. Influenza B Virus Victoria Group with a New Glycosylation Site Was Epidemic in Japan in the 2002-2003 Season. J Clin Microbiol 2004;42(7):3295–7.

9. Das SR, Hensley SE, David A, et al. Fitness costs limit influenza A virus hemagglutinin glycosylation as an immune evasion strategy. Proc Natl Acad Sci 2011;108(51):E1417–22.

10. Lee JM, Huddleston J, Doud MB, et al. Deep mutational scanning of hemagglutinin helps predict evolutionary fates of human H3N2 influenza variants. Proc Natl Acad Sci 2018;115(35):E8276–85.

11. Bae J-Y, Lee I, Kim JI, et al. A Single Amino Acid in the Polymerase Acidic Protein Determines the Pathogenicity of Influenza B Viruses. J Virol 2018;92(13):e00259–18.

12. Canaday LM, Resnick JD, Liu H, et al. HA and M2 sequences alter the replication of 2013–16 H1 live attenuated influenza vaccine infection in human nasal epithelial cell cultures. Vaccine 2022;40(32):4544–53.

13. Liu H, Grantham ML, Pekosz A. Mutations in the Influenza A Virus M1 Protein Enhance Virus Budding To Complement Lethal Mutations in the M2 Cytoplasmic Tail. J Virol 2018;92(1):e00858–17.

14. Wilson JL, Akin E, Zhou R, et al. The Influenza B Virus Victoria and Yamagata Lineages Display Distinct Cell Tropism and Infection-Induced Host Gene Expression in Human Nasal Epithelial Cell Cultures. Viruses 2023;15(9):1956.

15. Rowe T, Fletcher A, Lange M, et al. Delay of innate immune responses following influenza B virus infection affects the development of a robust antibody response in ferrets. mBio 2025;16(2):e02361–24.

16. Rowe T, Davis W, Wentworth DE, Ross T. Differential interferon responses to influenza A and B viruses in primary ferret respiratory epithelial cells. J Virol 2024;98(2):e01494–23.

17. Dudas G, Bedford T, Lycett S, Rambaut A. Reassortment between Influenza B Lineages and the Emergence of a Coadapted PB1–PB2–HA Gene Complex. Mol Biol Evol 2015;32(1):162–72.

18. McCullers JA, Wang GC, He S, Webster RG. Reassortment and Insertion-Deletion Are Strategies for the Evolution of Influenza B Viruses in Nature. J Virol 1999;73(9):7343–8.

19. Langat P, Raghwani J, Dudas G, et al. Genome-wide evolutionary dynamics of influenza B viruses on a global scale. PLOS Pathog 2017;13(12):e1006749.

20. Virk RK, Jayakumar J, Mendenhall IH, et al. Divergent evolutionary trajectories of influenza B viruses underlie their contemporaneous epidemic activity. Proc Natl Acad Sci 2020;117(1):619–28.

21. Sun X, Jayaraman A, Maniprasad P, et al. N-Linked Glycosylation of the Hemagglutinin Protein Influences Virulence and Antigenicity of the 1918 Pandemic and Seasonal H1N1 Influenza A Viruses. J Virol 2013;87(15):8756–66.

22. Sun W, Kang DS, Zheng A, et al. Antibody Responses toward the Major Antigenic Sites of Influenza B Virus Hemagglutinin in Mice, Ferrets, and Humans. J Virol 2019;93(2):e01673–18.

23. Rosu ME, Lexmond P, Bestebroer TM, et al. Substitutions near the HA receptor binding site explain the origin and major antigenic change of the B/Victoria and B/Yamagata lineages. Proc Natl Acad Sci 2022;119(42):e2211616119.

24. Page CK, Mubassir MHM, Chopra P, et al. N-glycosylation at the receptor binding site drives differences in receptor binding specificity between influenza B virus lineages. J Virol 2025;0(0):e01039–25.

25. Wilson JL, Zhou R, Liu H, Rothman R, Fenstermacher KZ, Pekosz A. Antigenic alteration of 2017-2018 season influenza B vaccine by egg-culture adaption. Front Virol [Internet] 2022 [cited 2023 Mar 5];2. Available from: https://www.frontiersin.org/articles/10.3389/fviro.2022.933440

26. Chen Z, Aspelund A, Jin H. Stabilizing the glycosylation pattern of influenza B hemagglutinin following adaptation to growth in eggs. Vaccine 2008;26(3):361–71.

27. Wilson JL, Zhou R, Liu H, Rothman R, Fenstermacher KZ, Pekosz A. Antigenic alteration of 2017-2018 season influenza B vaccine by egg-culture adaption. Front Virol [Internet] 2022 [cited 2025 May 9];2. Available from: https://www.frontiersin.org https://www.frontiersin.org/journals/virology/articles/10.3389/fviro.2022.933440/full

28. Propose new subclade of C.3 · influenza-clade-nomenclature/seasonal_B-Vic_HA@2afdc08 [Internet]. GitHub. [cited 2026 May 11];Available from: https://github.com/influenza-clade-nomenclature/seasonal_B-Vic_HA/commit/2afdc0867b7c2dbaba3d4ca5f9ad325cd2d36393

29. Brown SK, Tseng Y-Y, Aziz A, Baz M, Barr IG. Characterization of influenza B viruses with reduced susceptibility to influenza neuraminidase inhibitors. Antiviral Res 2022;200:105280.

30. Madsen A, Dai Y-N, McMahon M, et al. Human Antibodies Targeting Influenza B Virus Neuraminidase Active Site Are Broadly Protective. Immunity 2020;53(4):852–863.e7.

31. Liu H, Shaw-Saliba K, Westerbeck J, et al. Effect of human H3N2 influenza virus reassortment on influenza incidence and severity during the 2017–18 influenza season in the USA: a retrospective observational genomic analysis. Lancet Microbe [Internet] 2024 [cited 2024 July 20];0(0). Available from: https://www.thelancet.com/journals/lanmic/article/PIIS2666-5247(24)00067-3/fulltext

32. Vieira MC, Donato CM, Arevalo P, et al. Lineage-specific protection and immune imprinting shape the age distributions of influenza B cases. Nat Commun 2021;12(1):4313.

33. Edler P, Schwab LSU, Aban M, et al. Immune imprinting in early life shapes cross-reactivity to influenza B virus haemagglutinin. Nat Microbiol 2024;9(8):2073–83.

34. Shen J, Kirk BD, Ma J, Wang Q. Diversifying Selective Pressure on Influenza B Virus Hemagglutinin. J Med Virol 2009;81(1):114–24.

35. Nakagawa N, Kubota R, Maeda A, Okuno Y. Influenza B Virus Victoria Group with a New Glycosylation Site Was Epidemic in Japan in the 2002-2003 Season. J Clin Microbiol 2004;42(7):3295–7.

36. Gatherer D. Passage in egg culture is a major cause of apparent positive selection in influenza B hemagglutinin. J Med Virol 2010;82(1):123–7.

37. Recommended composition of influenza virus vaccines for use in the 2025-2026 northern hemisphere influenza season [Internet]. [cited 2025 July 11];Available from: https://www.who.int/publications/m/item/recommended-composition-of-influenza-virus-vaccines-for-use-in-the-2025-2026-nh-influenza-season

38. Recommended composition of influenza virus vaccines for use in the 2022-2023 northern hemisphere influenza season [Internet]. [cited 2025 June 30];Available from: https://www.who.int/publications/m/item/recommended-composition-of-influenza-virus-vaccines-for-use-in-the-2022-2023-northern-hemisphere-influenza-season

39. Recommended composition of influenza virus vaccines for use in the 2023-2024 northern hemisphere influenza season [Internet]. [cited 2025 June 23];Available from: https://www.who.int/publications/m/item/recommended-composition-of-influenza-virus-vaccines-for-use-in-the-2023-2024-northern-hemisphere-influenza-season

40. Saito T, Nakaya Y, Suzuki T, et al. Antigenic alteration of influenza B virus associated with loss of a glycosylation site due to host-cell adaptation. J Med Virol 2004;74(2):336–43.

41. Goldstein EJ, Harvey WT, Wilkie GS, et al. Integrating patient and whole-genome sequencing data to provide insights into the epidemiology of seasonal influenza A(H3N2) viruses. Microb Genomics 2018;4(1):e000137.

42. Madsen A, Dai Y-N, McMahon M, et al. Human Antibodies Targeting Influenza B Virus Neuraminidase Active Site Are Broadly Protective. Immunity 2020;53(4):852–863.e7.

43. Haghpanah F, Hamilton A, Klein E. Modeling the potential health impacts of delayed strain selection on influenza hospitalization and mortality with mRNA vaccines. Vaccine X 2023;14:100287.

44. Frutos AM, Cleary S, Reeves EL, et al. Interim Estimates of 2024-2025 Seasonal Influenza Vaccine Effectiveness - Four Vaccine Effectiveness Networks, United States, October 2024-February 2025. MMWR Morb Mortal Wkly Rep 2025;74(6):83–90.

45. Rose AM, Lucaccioni H, Marsh K, et al. Interim 2024/25 influenza vaccine effectiveness: eight European studies, September 2024 to January 2025. Eurosurveillance 2025;30(7):2500102.

46. National, Regional, and State Level Outpatient Illness and Viral Surveillance [Internet]. [cited 2025 July 8];Available from: https://gis.cdc.gov/grasp/fluview/fluportaldashboard.html

47. CDC. Weekly US Influenza Surveillance Report: Key Updates for Week 20, ending May 17, 2025 [Internet]. FluView. 2025 [cited 2025 July 8];Available from: https://www.cdc.gov/fluview/surveillance/2025-week-20.html

48. Ben Moussa M, Nwosu A, Schmidt K, et al. National Influenza Annual Report 2023–2024: A focus on influenza B and public health implications. Can Commun Dis Rep 50(11):393–9.

49. Zhou B, Lin X, Wang W, et al. Universal Influenza B Virus Genomic Amplification Facilitates Sequencing, Diagnostics, and Reverse Genetics. J Clin Microbiol 2014;52(5):1330–7.

50. Nogales A, Perez DR, Santos J, Finch C, Martfnez-Sobrido L. Reverse Genetics of Influenza B Viruses [Internet]. In: Perez DR, editor. Reverse Genetics of RNA Viruses: Methods and Protocols. New York, NY: Springer; 2017 [cited 2025 Jan 29]. p. 205–38.Available from: 10.1007/978-1-4939-6964-7_14

51. Cardenas-Garcia S, Caceres CJ, Rajao D, Perez DR. Reverse genetics for influenza B viruses and recent advances in vaccine development. Curr Opin Virol 2020;44:191–202.

52. Neher RA, Huddleston J, Bedford T, et al. Nomenclature for Tracking of Genetic Variation of Seasonal Influenza Viruses. Influenza Other Respir Viruses 2026;20(2):e70230.

53. Aksamentov I, Roemer C, Hodcroft EB, Neher RA. Nextclade: clade assignment, mutation calling and quality control for viral genomes. J Open Source Softw 2021;6(67):3773.

54. Hadfield J, Megill C, Bell SM, et al. Nextstrain: real-time tracking of pathogen evolution. Bioinformatics 2018;34(23):4121–3.

55. Tomasello G, Armenia I, Molla G. The Protein Imager: a full-featured online molecular viewer interface with server-side HQ-rendering capabilities. Bioinformatics 2020;36(9):2909–11.

56. Huddleston J, Hadfield J, Sibley TR, et al. Augur: a bioinformatics toolkit for phylogenetic analyses of human pathogens. J Open Source Softw 2021;6(57):2906.

57. Nguyen L-T, Schmidt HA, von Haeseler A, Minh BQ. IQ-TREE: A Fast and Effective Stochastic Algorithm for Estimating Maximum-Likelihood Phylogenies. Mol Biol Evol 2015;32(1):268–74.

58. Sagulenko P, Puller V, Neher RA. TreeTime: Maximum-likelihood phylodynamic analysis. Virus Evol 2018;4(1):vex042.

59. Engler JB. Tidyplots empowers life scientists with easy code-based data visualization. iMeta 2025;4(2):e70018.

