## supplemental tables and figures for "Emergence of an Antigenically Drifted and Reassorted Influenza B Virus at the end of the 2024-25 Influenza Season"

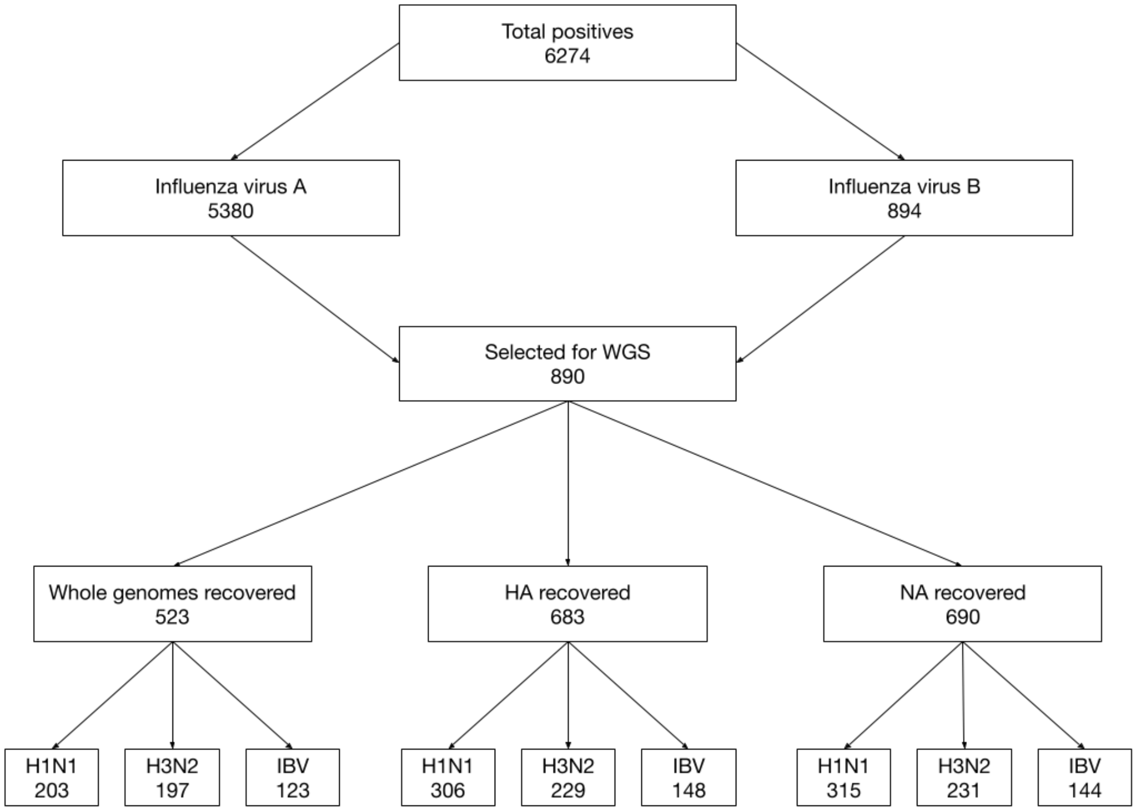


**Supplemental Figure S1.** Flow chart illustrating the number of samples selected for whole genome sequencing. HA and NA segments illustrate specimens where full-length HA segments were recovered.

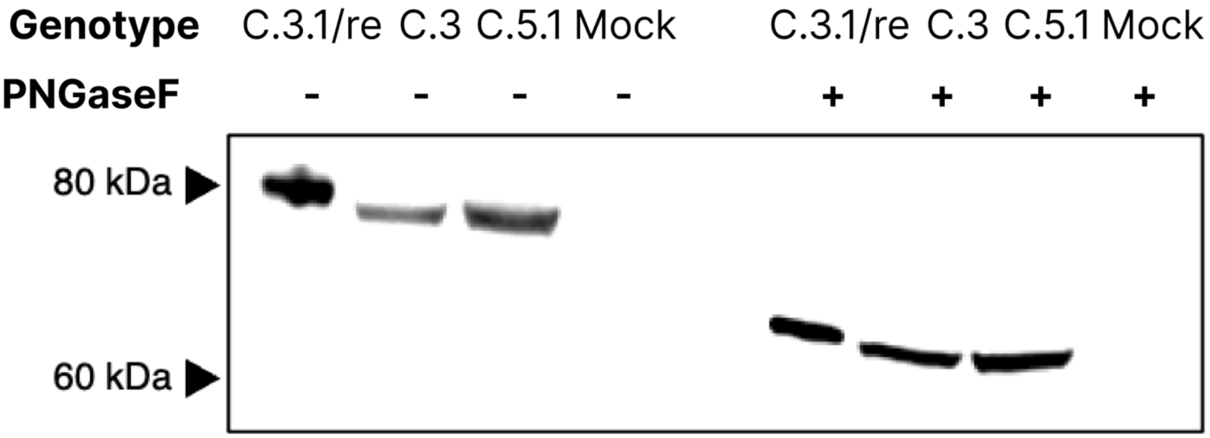


**Supplemental Figure S2 – The C.3re HA encodes additional N-linked glycan**. Denaturing SDS_PAGE gel followed by western blot of B HA. MDCK-SiaT-1 cells were infected with C.3re (B/Baltimore/JH-1192/2025), C.3 (B/Baltimore/JH-898/2023), or C.5.1 (B/Baltimore/JH-547/2024) and harvested after 48 hours at 33˚C. Samples were de-N-glycsoylated by PNGAseF treatment and blotted in parallel.

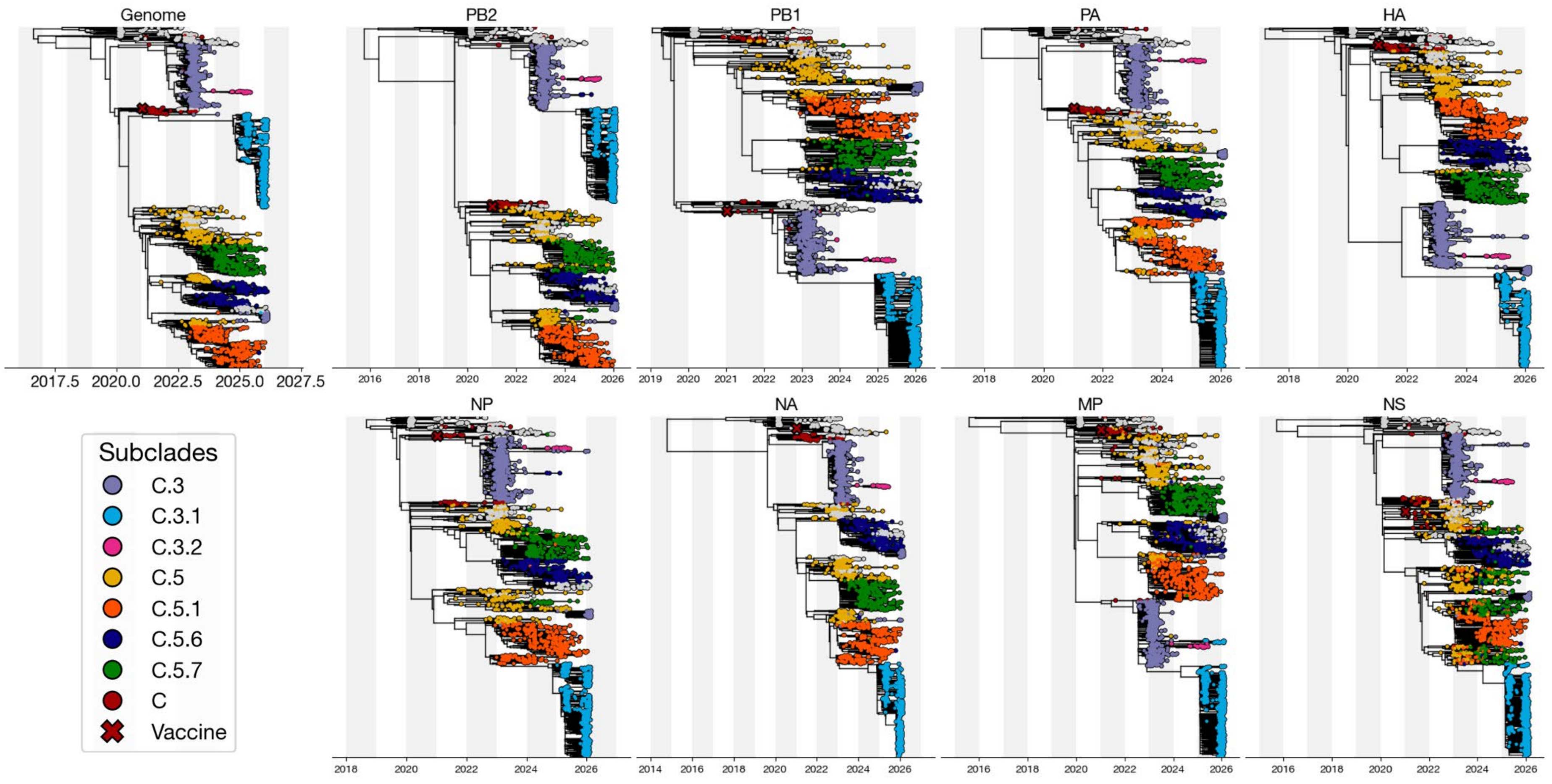


**Supplemental Figure S3 –** Gene tree phylogenies of the concatenated genome and 8 gene segments. The 2024-25 Influenza B vaccine component strain, B/Austria/1359417 /2021 is highlighted by an “X” on each phylogeny.

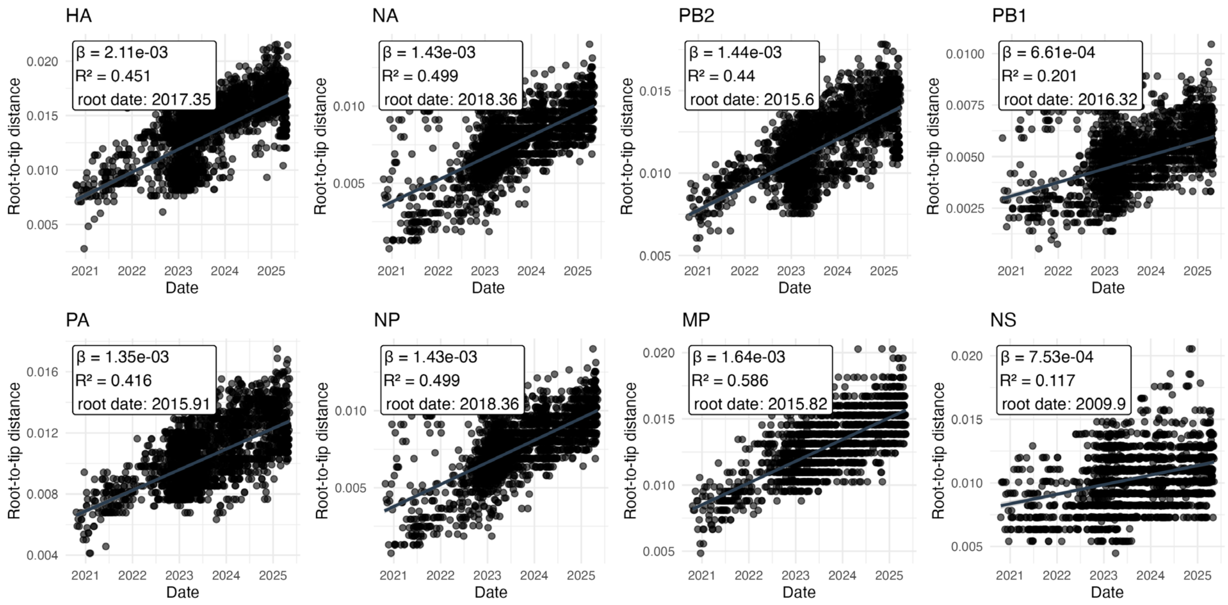


**Supplemental Figure S4 –** Root-to-tip regression of Influenza B segment phylogenies constructed in treetime dated to the calculated Y intercept (root date). β represents substitutions per site per year based on either collection or sequencing date.

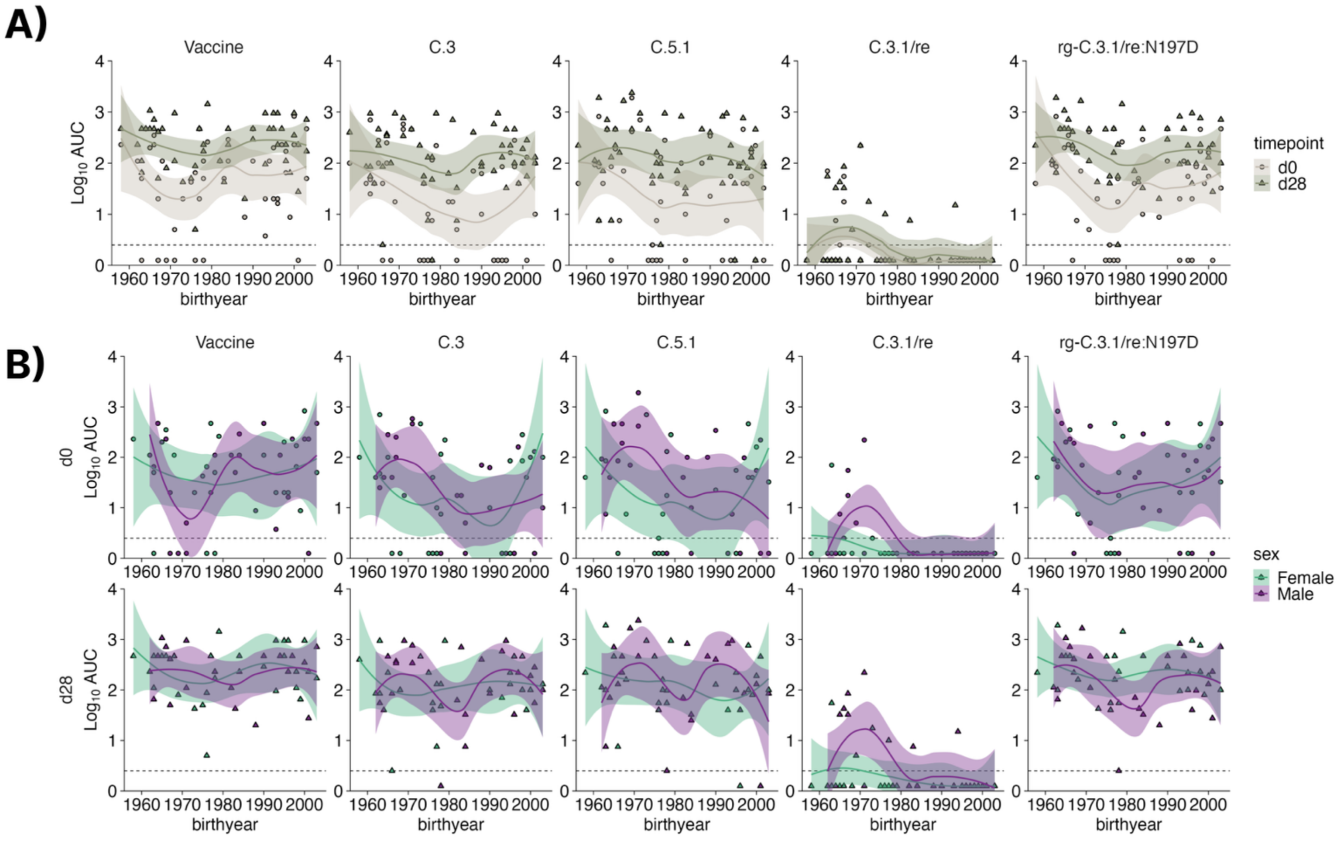


**Supplemental Figure S5. Serum neutralization area under the curve (AUC) values plotted against participant birth year. (A)** Locally Estimated Scatterplot Smoothing (LOESS) implemented in ggplot2 against baseline (d0) and post-vaccination (d28) neutralizing antibody AUC titers for the 2024-25 IBV vaccine (B/Austria/2021, C.3 (B/Baltimore/JH-898/2023), C.5.1 (B/Baltimore/JH-547/2024), C.3.1/re (B/Baltimore/JH-1192/2025) and rg-C.3.1/re:N197D (B/Baltimore/JH-1192/2025-background). The dotted line represents the limit of detection (AUC = 1.25). **(B)** AUC titers with LOESS fit by biological sex of participants. Statistical tests for age and sex are reported in Table 2. Only C.3.1/re was statistically significant (p=0.005; Kruskal-Wallis nonparametric test with Bonferroni's correction) by birthyear.

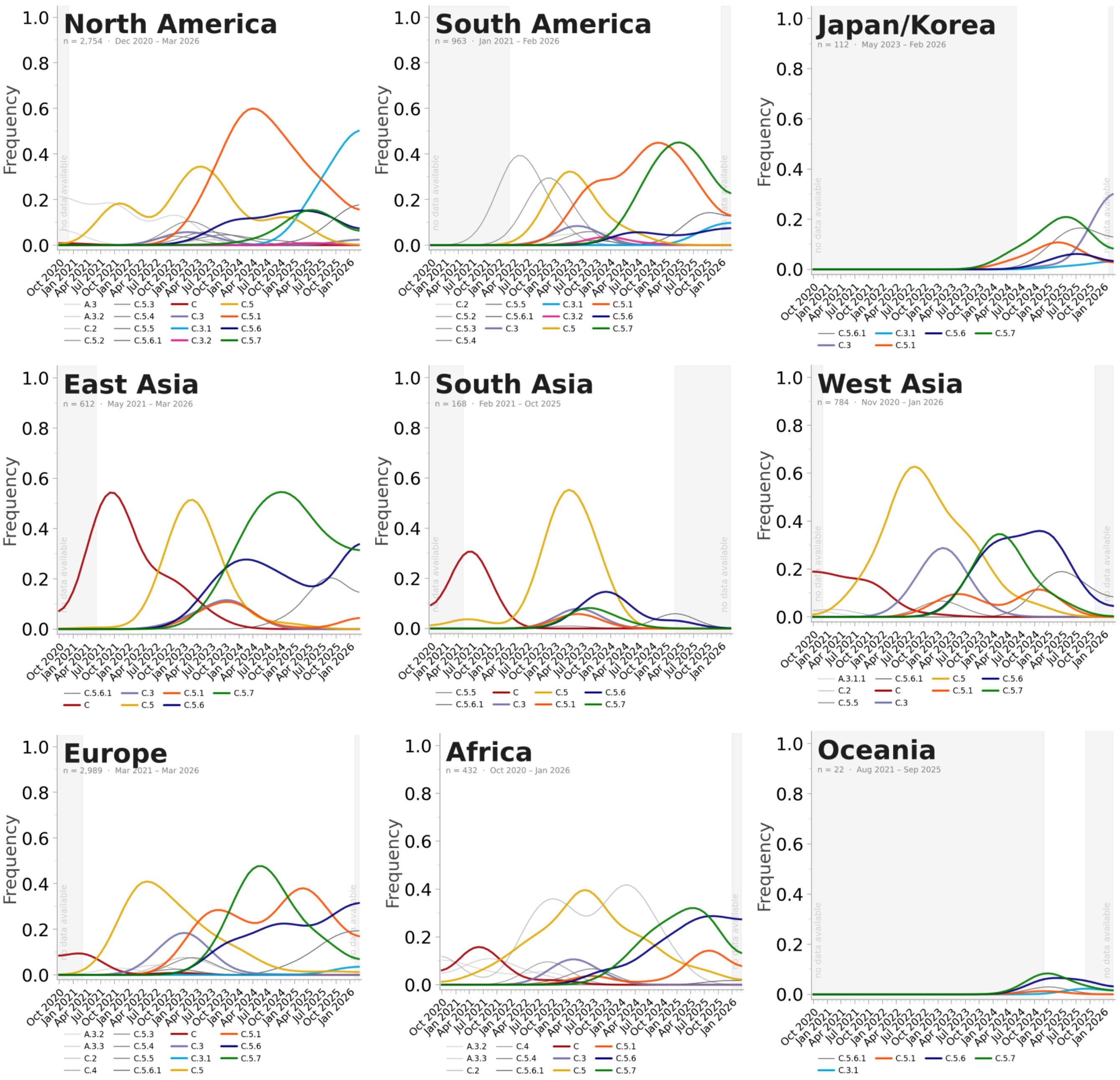


**Supplemental Figure S6.** Raw global influenza B subclade frequency between October 2020 and March 2026. Ingested sequence data are representative of all HA sequences available in GISAID between October 1, 2020 and March 18, 2026 to visualize relative frequency from all starting sequences. Sequence filtering and visualization scripts are available the Quarto notebook: https://github.com/elginakin/influenzab_c.3/blob/main/notebooks/06_frequencies.qmd

**Supplemental Table S1.** Total Influenza case counts in the Johns Hopkins Hospital in the 2024-25 season.

| Date | IAV Count | IBV Count | Total Cases | IAV % | IBV % |
|---:|---:|---:|---:|---:|---:|
| Oct 2024 | 2 | 0 | 2 | 100.0 | 0.0 |
| Nov 2024 | 57 | 4 | 61 | 93.4 | 6.6 |
| Dec 2024 | 510 | 12 | 522 | 97.7 | 2.3 |
| Jan 2025 | 1988 | 41 | 2029 | 98.0 | 2.0 |
| Feb 2025 | 2229 | 65 | 2294 | 97.2 | 2.8 |
| Mar 2025 | 526 | 372 | 898 | 58.6 | 41.4 |
| Apr 2025 | 53 | 400 | 453 | 11.7 | 88.3 |
| **TOTAL** | **5365** | **894** | **6259** | **85.7** | **14.3** |

**Supplemental Table S2.** Clinical and demographic characteristics of patients from the Johns Hopkins Hospital using a sub-cohort of infected individuals with recoverable high-quality IBV whole genome sequences.

| | Whole Cohort | | C.3 | | C.5.X | |
|---|---|---|---|---|---|---|
| | count | %N | count | %N | count | %N |
| No. of patients | 113 | | 67 | | 46 | |
| Female | 61 | 54.00% | 39 | 58.20% | 22 | 47.80% |
| Male | 52 | 46.00% | 28 | 41.80% | 24 | 52.20% |
| **Age Range** | | | | | | |
| Mean (SD) | 14 | 13 | 10.8 | 8.94 | 18.6 | 16.2 |
| Median | 9 | | 8 | | 13 | |
| Range | 57 | | 38 | | 57 | |
| <2 | 8 | 7.10% | 4 | 6.00% | 4 | 8.70% |
| 2-5 | 23 | 20.40% | 16 | 23.90% | 7 | 15.20% |
| 6-11 | 35 | 31.00% | 27 | 40.30% | 8 | 17.40% |
| 12-17 | 17 | 15.00% | 8 | 11.90% | 9 | 19.60% |
| 18-44 | 25 | 22.10% | 12 | 17.90% | 13 | 28.30% |
| 45-55 | 4 | 3.50% | 0 | 0.00% | 4 | 8.70% |
| 56-70 | 1 | 0.90% | 0 | 0.00% | 1 | 2.20% |
| **Comorbidities** | | | | | | |
| Hypertension | 18 | 15.90% | 9 | 13.40% | 9 | 19.60% |
| Lung Disease | 39 | 34.50% | 21 | 31.30% | 18 | 39.10% |
| Kidney Disease | 18 | 15.90% | 12 | 17.90% | 6 | 13.00% |
| Immunosupression | 24 | 21.20% | 11 | 16.40% | 13 | 28.30% |
| Diabetes | 9 | 8.00% | 3 | 4.50% | 6 | 13.00% |
| Heart failure | 6 | 5.30% | 2 | 3.00% | 4 | 8.70% |
| Smoker | 8 | 7.10% | 5 | 7.50% | 3 | 6.50% |
| Cancer | 34 | 30.10% | 23 | 34.30% | 11 | 23.90% |
| Pregnant | 1 | 0.90% | 0 | 0.00% | 1 | 2.20% |
| **Outcome** | | | | | | |
| Admitted | 20 | 17.70% | 12 | 17.90% | 8 | 17.40% |
| ICU | 4 | 3.50% | 2 | 3.00% | 2 | 4.30% |
| Supplemental Oxygen | 10 | 8.80% | 4 | 6.00% | 6 | 13.00% |
| Death | 1 | 0.90% | 0 | 0.00% | 1 | 2.20% |
| **Symptoms** | | | | | | |
| Chart count | 102 | | 59 | | 43 | |
| Fever | 32 | 31.40% | 21 | 35.60% | 11 | 25.60% |
| Cough | 10 | 9.80% | 10 | 16.90% | 4 | 9.30% |
| Headache | 2 | 2.00% | 1 | 1.70% | 1 | 2.30% |
| Breathing problems | 1 | 1.00% | 1 | 1.70% | 0 | 0.00% |
| Chest pain | 7 | 6.90% | 2 | 3.40% | 5 | 11.60% |
| Sore throat | 5 | 4.90% | 3 | 5.10% | 2 | 4.70% |
| URI | 6 | 5.90% | 4 | 6.80% | 2 | 4.70% |
| Abdominal pain | 9 | 8.80% | 5 | 8.50% | 4 | 9.30% |
| Emesis | 7 | 6.90% | 5 | 8.50% | 2 | 4.70% |
| Flu-like symptoms | 32 | 31.40% | 18 | 30.50% | 14 | 32.60% |
| Weakness | 1 | 1.00% | 1 | 1.70% | 0 | 0.00% |
| Seizures | 2 | 2.00% | 1 | 1.70% | 1 | 2.30% |

**Supplemental Table S3**. Viruses used in this study

| Strain Name | HA Clade | HA Subclade | Genotype | Total N-Linked Glycosylation Motifs | GISAID Accession |
|---|---|---|---|---|---|
| B/Baltimore/JH-1192/2025 | V1A.3a.2 | C.3.1 | C.3.1/re | 11 | EPI_ISL_19919784 |
| B/Baltimore/JH-547/2024 | V1A.3a.2 | C.5.1 | N/A | 10 | EPI_ISL_19861034 |
| B/Baltimore/JH-898/2024 | V1A.3a.2 | C.3 | N/A | 10 | EPI_ISL_20247894 |
| B/Austria/1359417/2021 | V1A.3a.2 | C | N/A | 10 | EPI_ISL_983345 |
| rg-B/Baltimore/JH-1192/2025:[N197D] | V1A.3a.2 | C.3.1 | N/A | 10 | |

**Supplemental Table S4.** JH-CEIRR influenza vaccine cohort age and sex study design

| Age Group (Years) | n(Female/Male) |
|---|---|
| 21-29 | 10 (5/5) |
| 30-39 | 10 (5/5) |
| 40-49 | 10 (5/5) |
| 50-59 | 10 (5/5) |
| 60-69 | 10 (5/5) |
| Total | 50 (25/25) |

**Supplemental Table S5.** JH-CEIRR influenza vaccine cohort with demographics, Neutralization Titer 50% and Area Under the Curve Values (AUC) by patient.

| Demographic Information | | | B/Victoria 2024-25 Vaccine B/Austria/1359417/2021-Egg | | | | B/Victoria C.3 B/Baltimore/JH-898/2023 | | | | B/Victoria C.5.1 B/Baltimore/JH-547/2024 | | | | B/Victoria C.3.1/re B/Baltimore/JH-1192/2025 | | | | B/Victoria rg-C.3.1/re:N197D B/Baltimore/JH-1192/2025 | | | |
|---|---|---|---|---|---|---|---|---|---|---|---|---|---|---|---|---|---|---|---|---|---|---|
| | | | Day 0 | | Day 28 | | Day 0 | | Day 28 | | Day 0 | | Day 28 | | Day 0 | | Day 28 | | Day 0 | | Day 28 | |
| Study ID | age | sex | NT50 | AUC | NT50 | AUC | NT50 | AUC | NT50 | AUC | NT50 | AUC | NT50 | AUC | NT50 | AUC | NT50 | AUC | NT50 | AUC | NT50 | AUC |
| 02-17-Pro-1208 | 23 | Male | 320 | 470 | 320 | 710 | 20 | 10 | 80 | 100 | 10 | 0 | 80 | 85 | 10 | 0 | 10 | 0 | 320 | 470 | 320 | 710 |
| 02-17-Pro-1231 | 23 | Female | 40 | 50 | 160 | 170 | 40 | 100 | 160 | 130 | 40 | 32.5 | 80 | 100 | 10 | 0 | 10 | 0 | 40 | 32.5 | 80 | 100 |
| 02-17-Pro-1190 | 25 | Male | 5 | 0 | 20 | 27.5 | 10 | 0 | 40 | 55 | 10 | 0 | 10 | 0 | 10 | 0 | 10 | 0 | 5 | 0 | 20 | 27.5 |
| 02-17-Pro-1206 | 25 | Male | 160 | 230 | 160 | 230 | 80 | 130 | 160 | 280 | 160 | 220 | 320 | 457.5 | 10 | 0 | 10 | 0 | 160 | 230 | 160 | 230 |
| 02-17-Pro-1228 | 26 | Female | 640 | 830 | 640 | 950 | 80 | 100 | 80 | 100 | 80 | 160 | 160 | 190 | 10 | 0 | 10 | 0 | 80 | 160 | 160 | 190 |
| 02-17-Pro-1236 | 26 | Male | 160 | 230 | 320 | 350 | 80 | 100 | 160 | 190 | 80 | 55 | 80 | 130 | 10 | 0 | 10 | 0 | 80 | 55 | 80 | 130 |
| 02-17-Pro-1268 | 27 | Female | 10 | 8.75 | 40 | 65 | 40 | 40 | 40 | 40 | 40 | 40 | 40 | 80 | 10 | 0 | 10 | 0 | 40 | 40 | 40 | 80 |
| 02-17-Pro-1260 | 28 | Female | 160 | 170 | 160 | 230 | 40 | 85 | 80 | 100 | 40 | 40 | 80 | 100 | 10 | 0 | 10 | 0 | 160 | 170 | 160 | 230 |
| 02-17-Pro-1265 | 28 | Female | 40 | 65 | 40 | 110 | 160 | 280 | 320 | 460 | 320 | 460 | 640 | 760 | 10 | 0 | 10 | 0 | 320 | 460 | 640 | 760 |
| 02-17-Pro-1234 | 29 | Male | 160 | 230 | 320 | 470 | 160 | 160 | 160 | 220 | 80 | 97.5 | 80 | 85 | 10 | 0 | 10 | 0 | 80 | 85 | 80 | 97.5 |
| 02-17-pro-1200 | 30 | Female | 20 | 20 | 640 | 950 | 10 | 0 | 320 | 460 | 10 | 0 | 320 | 0 | 10 | 0 | 10 | 0 | 20 | 20 | 640 | 950 |
| 02-17-Pro-1241 | 30 | Male | 20 | 16.25 | 160 | 230 | 10 | 0 | 80 | 32.5 | 10 | 0 | 40 | 40 | 10 | 0 | 10 | 0 | 10 | 0 | 40 | 40 |
| 02-17-Pro-1221 | 31 | Female | 160 | 200 | 320 | 470 | 80 | 85 | 80 | 130 | 40 | 55 | 80 | 100 | 10 | 0 | 10 | 0 | 40 | 55 | 160 | 100 |
| 02-17-Pro-1226 | 31 | Female | 20 | 20 | 640 | 950 | 10 | 0 | 160 | 220 | 10 | 0 | 160 | 220 | 10 | 0 | 10 | 0 | 10 | 0 | 160 | 220 |
| 02-17-Pro-1290 | 32 | Male | 80 | 110 | 320 | 470 | 10 | 0 | 640 | 940 | 20 | 7.5 | 640 | 940 | 10 | 0 | 20 | 15 | 80 | 110 | 320 | 470 |
| 02-17-Pro-1198 | 33 | Female | 20 | 20 | 80 | 95 | 10 | 0 | 80 | 70 | 10 | 0 | 40 | 40 | 10 | 0 | 10 | 0 | 20 | 20 | 320 | 95 |
| 02-17-Pro-1285 | 33 | Male | 5 | 3.75 | 640 | 950 | 10 | 0 | 320 | 580 | 10 | 0 | 640 | 820 | 10 | 0 | 10 | 0 | 10 | 0 | 640 | 820 |
| 02-17-Pro-1213 | 36 | Female | 80 | 110 | 160 | 290 | 20 | 10 | 160 | 100 | 40 | 22.5 | 80 | 127.5 | 10 | 0 | 10 | 0 | 80 | 110 | 160 | 290 |
| 02-17-Pro-1218 | 36 | Male | 320 | 470 | 320 | 470 | 80 | 62.5 | 80 | 85 | 160 | 340 | 320 | 400 | 10 | 0 | 10 | 0 | 320 | 470 | 320 | 470 |
| 02-17-Pro-1312 | 38 | Male | 10 | 8.75 | 20 | 20 | 80 | 70 | 320 | 400 | 80 | 100 | 320 | 460 | 10 | 0 | 10 | 0 | 10 | 8.75 | 20 | 20 |
| 02-17-Pro-1249 | 42 | Male | 80 | 110 | 160 | 230 | 20 | 5 | 40 | 7.5 | 20 | 10 | 40 | 32.5 | 10 | 0 | 10 | 0 | 20 | 10 | 40 | 32.5 |
| 02-17-Pro-1261 | 42 | Male | 160 | 290 | 320 | 470 | 10 | 0 | 40 | 32.5 | 10 | 0 | 40 | 25 | 10 | 0 | 10 | 0 | 160 | 290 | 320 | 470 |
| 02-17-Pro-1304 | 43 | Male | 40 | 50 | 40 | 42.5 | 20 | 17.5 | 640 | 700 | 80 | 100 | 640 | 820 | 10 | 0 | 20 | 7.5 | 40 | 50 | 40 | 42.5 |
| 02-17-Pro-1256 | 44 | Male | 80 | 110 | 80 | 110 | 80 | 17.5 | 80 | 62.5 | 40 | 40 | 80 | 70 | 10 | 0 | 10 | 0 | 40 | 40 | 80 | 70 |
| 02-17-Pro-1194 | 47 | Female | 160 | 260 | 1280 | 1430 | 160 | 220 | 640 | 940 | 160 | 280 | 640 | 940 | 10 | 0 | 10 | 0 | 160 | 260 | 1280 | 1430 |
| 02-17-pro-1203 | 48 | Female | 40 | 50 | 160 | 230 | 160 | 115 | 160 | 130 | 80 | 130 | 80 | 97.5 | 10 | 0 | 10 | 0 | 40 | 50 | 160 | 230 |
| 02-17-pro-1338 | 48 | Female | 5 | 1.25 | 160 | 170 | 20 | 7.5 | 40 | 47.5 | 20 | 17.5 | 40 | 55 | 10 | 0 | 10 | 0 | 20 | 17.5 | 40 | 55 |
| 02-17-Pro-1362 | 48 | Male | 80 | 110 | 160 | 230 | 10 | 0 | 10 | 0 | 10 | 0 | 10 | 2.5 | 10 | 0 | 10 | 0 | 10 | 0 | 10 | 2.5 |
| 02-17-Pro-1257 | 49 | Female | 20 | 20 | 320 | 470 | 20 | 10 | 160 | 130 | 10 | 0 | 160 | 160 | 10 | 0 | 10 | 0 | 10 | 0 | 160 | 160 |
| 02-17-Pro-1321 | 49 | Female | 320 | 470 | 320 | 470 | 10 | 0 | 20 | 7.5 | 20 | 7.5 | 80 | 100 | 10 | 0 | 20 | 10 | 320 | 470 | 320 | 470 |
| 02-17-pro-1277 | 50 | Female | 40 | 65 | 80 | 87.5 | 10 | 0 | 40 | 40 | 10 | 2.5 | 40 | 40 | 10 | 0 | 10 | 0 | 10 | 2.5 | 40 | 40 |
| 02-17-Pro-1359 | 50 | Female | 10 | 1.25 | 10 | 5 | 10 | 40 | 160 | 55 | 80 | 0 | 80 | 55 | 10 | 0 | 10 | 0 | 80 | 0 | 80 | 55 |
| 02-17-pro-1343 | 51 | Female | 40 | 42.5 | 40 | 50 | 10 | 0 | 160 | 220 | 10 | 0 | 320 | 460 | 10 | 0 | 10 | 0 | 10 | 0 | 320 | 460 |
| 02-17-Pro-1314 | 53 | Female | 20 | 20 | 40 | 42.5 | 320 | 460 | 320 | 460 | 640 | 700 | 640 | 940 | 10 | 2.5 | 20 | 17.5 | 20 | 20 | 40 | 42.5 |
| 02-17-Pro-1307 | 55 | Male | 10 | 5 | 80 | 110 | 320 | 580 | 640 | 760 | 1280 | 1900 | 1280 | 2380 | 160 | 220 | 160 | 220 | 10 | 5 | 80 | 110 |
| 02-17-pro-1374 | 55 | Male | 5 | 1.25 | 640 | 950 | 320 | 460 | 320 | 340 | 320 | 415 | 320 | 460 | 10 | 0 | 10 | 0 | 320 | 415 | 320 | 460 |
| 02-17-pro-1377 | 57 | Male | 5 | 1.25 | 40 | 80 | 10 | 17.5 | 640 | 940 | 80 | 70 | 1280 | 1660 | 20 | 5 | 20 | 5 | 80 | 70 | 1280 | 1660 |
| 02-17-Pro-1308 | 58 | Female | 80 | 110 | 320 | 470 | 10 | 0 | 80 | 100 | 20 | 7.5 | 160 | 160 | 10 | 0 | 10 | 0 | 20 | 7.5 | 160 | 160 |

| | | | | | | | | | | | | | | | | | | | | | | |
|---|---|---|---|---|---|---|---|---|---|---|---|---|---|---|---|---|---|---|---|---|---|---|
| 02-17-pro-1331 | 59 | Male | 20 | 20 | 40 | 50 | 160 | 250 | 160 | 370 | 160 | 190 | 160 | 220 | 40 | 55 | 80 | 85 | 160 | 190 | 160 | 220 |
| 02-17-Pro-1378 | 59 | Male | 5 | 1.25 | 320 | 410 | 80 | 100 | 320 | 340 | 320 | 460 | 320 | 460 | 20 | 17.5 | 40 | 32.5 | 5 | 1.25 | 320 | 410 |
| 02-17-Pro-1201 | 60 | Male | 160 | 230 | 640 | 710 | 40 | 40 | 80 | 85 | 80 | 85 | 80 | 130 | 10 | 2.5 | 40 | 42.5 | 160 | 230 | 640 | 710 |
| 02-17-Pro-1313 | 60 | Female | 160 | 350 | 320 | 470 | 10 | 0 | 10 | 2.5 | 10 | 0 | 20 | 7.5 | 10 | 0 | 10 | 0 | 160 | 350 | 320 | 470 |
| 02-17-Pro-1259 | 61 | Female | 160 | 200 | 320 | 470 | 80 | 100 | 160 | 220 | 320 | 452.5 | 320 | 460 | 10 | 0 | 10 | 0 | 160 | 200 | 320 | 470 |
| 02-17-Pro-1262 | 61 | Male | 160 | 200 | 320 | 1070 | 320 | 280 | 320 | 460 | 320 | 460 | 640 | 700 | 20 | 7.5 | 40 | 32.5 | 160 | 200 | 320 | 1070 |
| 02-17-Pro-1305 | 62 | Male | 320 | 470 | 320 | 470 | 40 | 40 | 40 | 40 | 40 | 40 | 80 | 70 | 10 | 0 | 10 | 0 | 320 | 470 | 320 | 470 |
| 02-17-Pro-1185 | 63 | Male | 40 | 65 | 40 | 65 | 40 | 25 | 80 | 55 | 20 | 7.5 | 20 | 7.5 | 10 | 0 | 10 | 0 | 40 | 65 | 40 | 65 |
| 02-17-Pro-1281 | 63 | Female | 5 | 0 | 320 | 470 | 40 | 47.5 | 80 | 85 | 80 | 85 | 80 | 100 | 10 | 0 | 10 | 0 | 80 | 85 | 80 | 100 |
| 02-17-Pro-1323 | 63 | Female | 40 | 50 | 80 | 110 | 640 | 700 | 640 | 940 | 640 | 820 | 1280 | 1900 | 40 | 70 | 40 | 55 | 320 | 820 | 640 | 1900 |
| 02-17-pro-1324 | 64 | Male | 80 | 110 | 160 | 230 | 40 | 40 | 80 | 85 | 80 | 92.5 | 80 | 112.5 | 10 | 0 | 10 | 0 | 80 | 92.5 | 80 | 112.5 |
| 02-17-Pro-1186 | 68 | Female | 160 | 230 | 320 | 470 | 320 | 100 | 640 | 400 | 40 | 40 | 160 | 220 | 10 | 0 | 10 | 0 | 40 | 40 | 160 | 220 |
