## supplemental methods for "Emergence of an Antigenically Drifted and Reassorted Influenza B Virus at the end of the 2024-25 Influenza Season"

**Nucleic Acid Extraction and Whole Genome Amplification from clinical specimens**

Nucleic acid was extracted using the Chemagic Viral RNA/DNA Kit following the manufacturer’s instructions (Revvity, Waltham, MA, USA). The whole genomes of IBV were amplified using Invitrogen Supercript III (Waltham, MA, USA) and universal IBV primer cocktail.^1^ Library preparation was performed with NEBNext Artic SARS-CoV-2 Companion Kit (New England Biolabs, Ipswich, MA, USA), and sequencing was performed following manufacturer’s instructions, using R10.4.1 flow cells on a GridION (Oxford Nanopore Technologies, Oxford, UK).

**Influenza Genome Assembly**

Fastq files were demultiplexed using the artic_guppyplex tool (Artic version 1.2.2). Nucleotide sequence assembly was performed using the default settings of the FLU module of the Iterative Refinement Meta-Assembler (IRMA version 1.0.2) which include a minimum average quality score of 24 and a site depth of 100. The alignment of genomes and reference sequences, downloaded from GISAID, was performed using the built-in alignment tool in Nextclade.

Quality control scores for sequences were assigned using the built-in pipeline available in Nextclade. Sequences with overall quality scores of 30 and above were excluded from sequence analysis.

**Cell Culture**

Madin-Darby canine kidney (MDCK) cells (provided by Dr. Robert A. Lamb, Northwestern University), MDCK-SIAT-1 cells (provided by Dr. Scott Hensley, University of Pennsylvania) and MDCK-SIAT-1-TMPRSSII cells (provided by Dr. Jesse Bloom, Fred Hutchinson Cancer Center) were maintained in complete medium (CM) consisting of Dulbecco’s Modified Eagle Medium (DMEM) supplemented with 10% fetal bovine serum, 100 units/mL penicillin/streptomycin (Life Technologies) and 2 mM Glutamax (Gibco) at 37 °C and 5% CO2. Human nasal epithelial cells (hNEC) (PromoCell) were cultivated as previously described^2^.

**Serum Neutralization Assay (NT_50_)**

Serum samples used for this analysis originated from JHHS healthcare workers recruited from the Johns Hopkins Center of Excellence for Influenza Research and Response (JH-CEIRR) during the Fall 2024 influenza vaccination campaign. Serum samples collected at the time of vaccination or 28 days post vaccination were first treated (1:3 ratio serum to enzyme) with Receptor Destroying Enzyme (Denka-Seiken, Tokyo, Japan) and incubated overnight at 37 °C followed by inactivation at 57 °C for 35 min. Serum was diluted 2-fold in IM (Dulbecco modified Eagle medium (Sigma), with 10% penicillin/streptomycin (Gibco), 10% L-glutamine (Gibco), 0.5% Bovine Serum Albumin (Sigma), and 2.5 µg/mL of N-acetyl trypsin (Sigma) at 37 °C and 5% CO_2_) and 100 TCID50 was added for a one-hour incubation at room temperature. Serum Sample/Virus was used to infect a confluent layer of MDCK-SIAT-1cells. The inoculums were removed after 24 hours, IM supplemented with 2.5µg/mL N-acetyl Trypsin was replaced, and cells were incubated for 96 hours at 37°C. Plates were fixed and stained as described previously^3^. The Neutralizing antibody titer was calculated using the highest serum dilution that led to greater or equal to 50% cytopathic effect (CPE).

**Assessment of** **NA activity and oseltamivir sensitivity**

Influenza Neuraminidase Inhibitor Resistance Detection Kit (Thermofisher) was used per manufacturer’s protocol to investigate relative susceptibility of B/Baltimore/JH-1192/2025 and B/Baltimore/JH-547/2024 to neuraminidase-inhibitor Oseltamivir (oseltamivir carboxylate, MedChemExpress). Briefly, all viruses tested were serially diluted two-fold 11 times in NA-Star assay buffer and added to opaque white 96-well plates (Falcon). NA-Star assay substrate was diluted in NA-Star assay buffer and added to all wells per the manufacturer’s protocol, and plates were incubated at room temperature for 1 hour on a plate shaker. Plates were then removed from shaker, and immediately prior to reading each plate, NA-Star accelerator was added in equal volume to each well. Luminescence was measured on a SpectraMax I3 multimode plate reader (Agilent) and analyzed (Graphpad Prism 10). The amount of virus added was calculated as the dilution of virus that yielded 10^4^ Relative luciferase units (RLU) with minimal variability and a high signal-to-noise ratio (i.e., greater than 1000). Each virus was titrated in quadruplicate.

Oseltamivir (oseltamivir carboxylate, MedChemExpress) was serially diluted four-fold, starting at 50 µM. Virus was diluted to the selected input based on the titration output and added to the oseltamivir dilution series shaken briefly, and incubated at 37ºC/5% CO_2_ for 30 minutes. NA-Star assay substrate, diluted in NA-Star assay buffer was then added to each well, and plates were incubated for an additional 30 minutes at room temperature with vigorous shaking. NA-Star accelerator was added immediately prior to reading each plate. RLU quantified by the oseltamivir+virus mixture were normalized to wells with no virus (representing 100% NA inhibition) and to wells with only virus and no oseltamivir (representing 0% NA inhibition). Curves were fit to a nonlinear regression log(agonist) versus normalized response model, and IC_50_ values for oseltamivir-mediated NAI were calculated as 4.73 nM for B/Baltimore/JH-547/2024, and 3.53 nM for B/Baltimore/JH-1192/2025 (Graphpad Prism 10).

**Virus Isolation on Human Nasal Epithelial Cells (hNECs) and MDCK-SIAT-1-TMPRSSII cells**

Nasopharyngeal swabs or nasal washes from individuals who were influenza B positive were used for virus isolation on MDCK-SIAT-TMPRSS2 cells or human nasal epithelial cell (hNEC) cultures as described previously ^2,4^. The MDCK-SIAT-TMPRSS2 cells were cultured at 37°C with 5% CO2 in complete medium (CM) consisting of Dulbecco’s Modified Eagle Medium (DMEM) supplemented with 10% fetal bovine serum, 100 units/mL penicillin/streptomycin (Life Technologies, Carlsbad, CA, USA) and 2 mM Glutamax (Gibco, Waltham, MA, USA). For virus isolation, the cells were grown in 96 well dishes, washed twice with 300ul of phosphate buffered saline (PBS) and 50-90 µl of specimen was added to the cells and incubated for two hours at 37°C. The supernatant was then aspirated and cells were washed twice with 300 µl of PBS, then incubated at 33°C for three, five, and seven days in 100 µl of infection media (DMEM supplemented with 0.3% BSA (Sigma-Aldrich, St. Louis, MO, USA), 100 units/mL pen/strep, 2 mM Glutamax (Thermo-Fisher, Waltham, MA, USA), and 2.5 µg/mL of N-acetylated Trypsin from bovine pancreas (Millipore Sigma, Burlington, MA). TCID50 was performed on collected supernatants and virus seed stocks were made from the collected media when virus was detected at concentrations greater than 10^4^ TCID50/mL.

**Virus Cloning and Plasmids**

We used cDNA derived from virus seed stocks as the templates for PCR

The cDNA Influenza B C.3.1/re strain B/Baltimore/JH-1192/2025 was amplified by PCR with primers and cloned by either restriction ligation of HiFi assembly into the bi-directional plasmid pDP2002 (a gift from Dr. Daniel Perez) as previously described.^5^ Both the plasmid and individual segments were digested separately with restriction enzymes (either BsmBI or BsaI, New England BioLabs) and then combined and ligated together with T4 DNA ligase (New England BioLabs). Ligated plasmids were transformed into DH5α bacteria (Thermo Fisher) and grown overnight on LB plates containing carbenicillin to select for colonies that successfully received the plasmid. Individual colonies were isolated and grown, and plasmid was extracted using a Qiagen QIAprep Spin Miniprep Kit. Plasmid sequences were verified using restriction enzyme digestion and whole plasmid sequencing, which was performed by Plasmidsaurus using Oxford Nanopore Technology with custom analysis and annotation. Bacteria expressing the correct plasmid were grown in bulk and high-purity plasmid was extracted using ZymoPURE Plasmid Midiprep kit (Zymo research) and verified by sequencing of the coding region of the influenza virus segment. For generating a C.3.1/re HA lacking a NLG motif at residue 197, mutagenic primers were designed using the New England Biolabs online software NEBaseChanger PrimerF (5’-CCACTCTGACgatGAGACCCAAATG-3’) Primer R (5’- AACCCCCAAACGGTAATTTG-3’) The Q5 Site-Directed Mutagenesis Kit (New England BioLabs) was then used with the B/Baltimore/JH-1192/2025 hemagglutinin plasmid to produce plasmids with the desired mutations according to the manufacture instructions (NEB Catalog #E055DS). Plasmids were transformed, extracted, and confirmed as described above.

**Recombinant Virus Production**

Recombinant viruses were generated using the pDP2002 8 plasmid reverse genetic system previously described.^5,6^

**Virus Stock Preparation**

A T75 flask of confluent MDCK-SIAT-1 cells was infected at an MOI of 0.01 in IM supplemented with 2.5µg/mL of NAT. Working stocks for each vaccine strain or clinical isolate were generated by infecting a T75 flask of MDCK-SIAT-1 cells at an MOI of 0.001 for one hour at room temperature while rocking. The inoculum was removed, and cells were placed in a 33 °C incubator and monitored daily for CPE. Working stock was harvested between 3 and 5 days later, when CPE was seen in 75–80% of the cells. Harvested media were centrifuged at 400× *g* for 10 min at 8 °C to remove cell debris, and the resulting supernatant was aliquoted into 500 µL and stored at −80 °C. Infectious virus quantity of working stocks was determined using TCID50 assay.

**Tissue Culture Infectious Dose 50 (TCID50) Assay**

MDCK-SIAT-1 cells were seeded in a 96-well plate 2 days before assay and grown to 100% confluence. Cells were washed twice with PBS+ then 180 µL of IM was added to each well. Ten-fold serial dilutions of virus from 10^−1^ to 10^−7^ were created and then 20 µL of the virus dilution was added to the MDCK-SIAT-1 cells. Cells were incubated for 6 days at 33 °C then fixed with 2% formaldehyde. After fixing, cells were stained with Naphthol Blue Black, washed and virus titer was calculated using the Reed and Muench method.

**Validating hemagglutinin N-linked glycosylation motifs by PNGAseF treatment and western blot**

MDCK-SIAT-1 cells were seeded in 6-well plates and allowed to reach confluency. Twenty-four hours later, cells were infected at a multiplicity of infection (MOI) of 1 in infection medium supplemented with 2.5 µg/mL NAT. At 24 hours post-infection, cell culture supernatants were collected, and monolayers were washed once with PBS and lysed in 100 µL of 1× RIPA buffer supplemented with 1× cOmplete™ protease inhibitor cocktail (Roche) for 20 minutes on ice. Lysates were transferred to microcentrifuge tubes and clarified by centrifugation at 13,000 × g for 10 minutes at 4 °C. The clarified lysate was divided into PNGase F–treated and untreated aliquots.

PNGase F digestion (Promega, catalog #V483A) was performed under denaturing and reducing conditions using a final reaction mixture containing 0.25% SDS, 50 mM dithiothreitol (DTT), 50 mM sodium phosphate buffer (pH 7.5), and 1% NP-40. Digested and undigested lysates were analyzed by reducing SDS-PAGE, followed by dry transfer to PVDF membranes and immunoblotting with an anti-Influenza B HA monoclonal antibody (ThermoFisher MA5-29901, clone 004; rabbit monoclonal, 1:2000) following blocking in LI-COR Intercept™ protein-free blocking buffer (927-80001). Bound HA antigen was detected with an HRP-conjugated anti-rabbit secondary antibody (1:10,000 in blocking buffer). Chemiluminescence was developed using ECL substrate according to the manufacturer’s instructions and imaged immediately on an Azure 600 imaging system.

**Influenza B clade and genotype definitions**

Influenza B viruses were classified using algorithmic clade proposals for the HA segment implemented into Nextclade (last accessed March 18, 2026).^7–9^ As of March 18, 2026, the recognized C.3 subclades and their clade defining amino acid mutations include C.3 (128K, 154E), C.3.1 (197N, 208S), and C.3.2 (197N, 208P) available at: https://github.com/influenza-clade-nomenclature/seasonal_B-Vic_HA/tree/main/subclades.

**Influenza B Reassortment Nomenclature**

Reassortment events were identified through comparative phylogenetic analysis of all eight influenza B virus genomic segments. Viruses were classified as reassortants when incongruent phylogenetic placement was observed across segments, specifically where the hemagglutinin (HA) segment clustered within the C.3 or C.3.1 clade, while one or more non-HA segments clustered within the C.5 lineage. To facilitate reassortment events within this study, we implemented a nomenclature system based on the HA clade designation coupled with the time of discovery within the dataset. Reassortant genotypes were annotated using the format: <HA clade>/re[.X]. where <HA clade> denotes the HA phylogenetic clade, /re indicates that the virus represents a reassortant genotype, and .X is an optional numeric suffix used to distinguish multiple independently identified reassortment events within the same HA clade. Numbering of reassortment events (e.g., /re, /re.1) reflects the order of discovery within the dataset. At the time of initial detection (April 2025), HA clade assignments were based on the contemporaneous phylogenetic framework. As HA clade definitions were refined with the inclusion of additional sequence data, some reassortant lineages were subsequently reclassified. For example, viruses initially designated as C.3/re were later reassigned to C.3.1/re following recognition of a distinct monophyletic subclade (C.3.1) within the original C.3 grouping. In such cases, reassortment nomenclature was updated to reflect the revised HA clade designation while retaining the original reassortment classification. This nomenclature system was designed to provide an intuitive and flexible framework for tracking reassortant viruses within the context of the current HA clade classifications. However, because reassortment designations are directly tied to HA clade names, updates to HA phylogeny may necessitate retrospective renaming of reassortant lineages.

**Protein Structure modeling**

Mutations in Influenza B **Hemagglutinin** (PDB 4FQM) and Neuraminidase (PDB 4CPL) were visualized using Protein Imager ^10^ .

**Phylogenetic Reconstruction of C.3.1/re ancestry using Maximum Likelihood**

19,689 Influenza B genomes were access from GISAID filtered by original passage and collection date between October 2020 and August 22, 2025. 3836 whole genomes were randomly down sampled using scripts available in [01_ingest.qmd](https://github.com/elginakin/influenzab_c.3/blob/main/notebooks/01_ingest.qmd). In order to maximize temporal and geographic diversity of uploaded d genomes, we grouped down selection by collection week and country implemented in augur filter^11^. Concatenated genome- and gene-level phylogenetics (totaling to 9) were constructed using IQ-Tree2 available in the augur tree module ^23,24^. Observed NS segment polytomies were resolved using IQ-Tree2 with the following flags: -B 1000 -alrt 1000 -T 4 (S FFig. S3). Resulting phylogeny branches were refined using the GISAID-submitted collection date or Johns Hopkins Hospital Sequencing date using treetime implemented in augur refine^11,13^. Nextclade was used to call clade and subclade designations (<https://github.com/influenza-clade-nomenclature/seasonal_B-Vic_HA/blob/main/subclades/C.3.yml>; last accessed July 6^th^, 2025) in addition to quality metrics and putative glycosylation sites^8^

**Statistical analyses**

Statistical analyses were performed in R (4.3.2) using Tidyplots ^14^ or Graphpad Prism (10.4.2) and described in Fig. legends Time to the most recent common ancestor estimates and confidence intervals were performed using baltic^15^.

**Supplemental Methods References**

1.	Zhou, B. *et al.* Universal Influenza B Virus Genomic Amplification Facilitates Sequencing, Diagnostics, and Reverse Genetics. *J. Clin. Microbiol.* **52**, 1330–1337 (2014).

2.	Swanson, N. J. *et al.* 2019–2020 H1N1 clade A5a.1 viruses have better in vitro fitness compared with the co-circulating A5a.2 clade. *Sci. Rep.* **13**, 10223 (2023).

3.	Wilson, J. L. *et al.* The Influenza B Virus Victoria and Yamagata Lineages Display Distinct Cell Tropism and Infection-Induced Host Gene Expression in Human Nasal Epithelial Cell Cultures. *Viruses* **15**, 1956 (2023).

4.	Lee, J. M. *et al.* Deep mutational scanning of hemagglutinin helps predict evolutionary fates of human H3N2 influenza variants. *Proc. Natl. Acad. Sci.* **115**, E8276–E8285 (2018).

5.	Nogales, A., Perez, D. R., Santos, J., Finch, C. & Martínez-Sobrido, L. Reverse Genetics of Influenza B Viruses. in *Reverse Genetics of RNA Viruses: Methods and Protocols* (ed. Perez, D. R.) 205–238 (Springer, New York, NY, 2017). doi:10.1007/978-1-4939-6964-7_14.

6.	Cardenas-Garcia, S., Caceres, C. J., Rajao, D. & Perez, D. R. Reverse genetics for influenza B viruses and recent advances in vaccine development. *Curr. Opin. Virol.* **44**, 191–202 (2020).

7.	Neher, R. A. *et al.* Nomenclature for Tracking of Genetic Variation of Seasonal Influenza Viruses. *Influenza Other Respir. Viruses* **20**, e70230 (2026).

8.	Aksamentov, I., Roemer, C., Hodcroft, E. B. & Neher, R. A. Nextclade: clade assignment, mutation calling and quality control for viral genomes. *J. Open Source Softw.* **6**, 3773 (2021).

9.	Hadfield, J. *et al.* Nextstrain: real-time tracking of pathogen evolution. *Bioinformatics* **34**, 4121–4123 (2018).

10.	Tomasello, G., Armenia, I. & Molla, G. The Protein Imager: a full-featured online molecular viewer interface with server-side HQ-rendering capabilities. *Bioinformatics* **36**, 2909–2911 (2020).

11.	Huddleston, J. *et al.* Augur: a bioinformatics toolkit for phylogenetic analyses of human pathogens. *J. Open Source Softw.* **6**, 2906 (2021).

12.	Nguyen, L.-T., Schmidt, H. A., von Haeseler, A. & Minh, B. Q. IQ-TREE: A Fast and Effective Stochastic Algorithm for Estimating Maximum-Likelihood Phylogenies. *Mol. Biol. Evol.* **32**, 268–274 (2015).

13.	Sagulenko, P., Puller, V. & Neher, R. A. TreeTime: Maximum-likelihood phylodynamic analysis. *Virus Evol.* **4**, vex042 (2018).

14.	Engler, J. B. Tidyplots empowers life scientists with easy code-based data visualization. *iMeta* **4**, e70018 (2025).

15.	Dudas, G., Bedford, T., Lycett, S. & Rambaut, A. Reassortment between Influenza B Lineages and the Emergence of a Coadapted PB1–PB2–HA Gene Complex. *Mol. Biol. Evol.* **32**, 162–172 (2015).
